# Structures of native SV2A reveal the binding mode for tetanus neurotoxin and anti-epileptic racetams

**DOI:** 10.1101/2024.05.07.592960

**Authors:** Stephan Schenck, Toon Laeremans, Jan Steyaert, Janine D. Brunner

## Abstract

The synaptic vesicle glycoprotein 2A (SV2A) is a *bona fide* synaptic vesicle (SV) constituent of controversial function with homology to the major facilitator superfamily (MFS) and essential in vertebrate neurotransmission. Despite its high medical relevance as the target of the anti-epileptic drug Levetiracetam (LEV) and as receptor for clostridial neurotoxins (CNTs), among them several botulinum neurotoxin (BoNT) serotypes and potentially tetanus neurotoxin (TeNT), we lack detailed insight about these molecular interactions. We purified native SV2A from brain and subjected it to a structural analysis to advance our understanding of drug-binding to this enigmatic protein and explore structurally uncharacterized toxin-SV2A interfaces. Our analysis uncovered that TeNT binds SV2 proteins strikingly different from BoNT/A and delivers visual evidence for the dual receptor hypothesis through structurally resolved, co-purified gangliosides in the complex. The structures provide compelling support for SV2A as the protein receptor for TeNT in central neurons, recapitulate the geometry of CNT binding to dipartite SV2-ganglioside receptors on neuronal surfaces in a membrane-like constellation and have implications for toxin-engineering through a previously unknown SV2A-toxin interface. Further, a LEV-bound structure of SV2A reveals the drug-interacting residues and delineates a putative substrate pocket in SV2A. Our work provides an explanation for the SV2-isoform-specificity of LEV and its derivatives and paves the way for improved design of anti-convulsant drugs in epilepsy treatment.

## Introduction

Synaptic vesicles (SVs) that store neurotransmitters for release into the synaptic cleft upon fusion with the presynaptic membrane are among the best studied organelles in the cell^1^. Yet, several SV-residents have resisted efforts to unveil their precise function. Among those proteins is the synaptic vesicle glycoprotein 2 family (SV2), consisting of three members, SV2A-C in vertebrate genomes^2,3^. SV2A, cloned more than 30 years ago^4,5^, is a multipass membrane protein with MFS architecture and strict SV-localization and the most widely expressed isoform in the brain where it is found in inhibitory and excitatory neurons. It was early proposed to constitute a transporter due to substantial homology to sugar transporters and was further considered as a regulator of presynaptic Ca^2+^ levels^6^. However, a substrate, potentially even a neurotransmitter or modulator, could not be assigned to SV2A (but see^7^) and a Ca^2+^ uptake activity could not be supported^8^. The deletion of SV2A in mice leads to severe seizures resulting in fatality within three weeks after birth, underpinning its importance in the brain^6,9^. On electrophysiological level, the deletion of SV2A shows reduced evoked synaptic response but basic transmission parameters like miniature frequency and amplitude as well as the size of the readily releasable pool are unaffected^10^. SV2A seems to be required in a last step of the exocytotic cycle before Ca^2+^-elicited fusion of SVs, after docking and priming^2,10^. Rescue experiments in SV2A/B knock-out neurons have shown that conserved residues in the transmembrane moiety (Trp300 and Trp666) are required for SV2A function^11^. As with the early conjectures about SV2 proteins as transporters, neither a transported substance nor another molecular explanation could be suggested from these experiments. Nevertheless, the data is in support of a transport function because the conserved residues that have been mutated are transport-related in other MFS transporters.

SV2 proteins are interacting with the Ca^2+^ sensor synaptotagmin (Syt)^12–14^ during SV-retrieval and it has been suggested that reduced Syt-levels in SVs are causative for the observed deficits in SV2A deletion studies^15–17^. However, it was also reported that N-terminal deletion mutants of SV2A, eliminating the region that interacts with Syt, could rescue the SV2A/B DKO^10^. SV2 proteins were further suggested to display charged sugar moieties (keratan sulfates) as a matrix to increase transmitter storage^18^, but these structures have not been proven yet to be a common feature across vertebrate SV2 proteins. Many of the suggested functions of SV2 proteins are not exclusive, suggesting that these proteins fulfil multiple roles^2,3^.

Despite these many uncertainties regarding their physiological role, SV2 proteins are of high medical relevance. SV2A has been identified as the target of the major anti-epileptic drug Levetiracetam (LEV)^19,20^ that was FDA-approved in 1999 and became one of the most prescribed anti-seizure medications. Since the SV2A deletion results in severe seizures, an inhibitory action on the function of SV2A by LEV appears counterintuitive. However, a mutagenesis scan of SV2A using radiolabelled racetams in binding assays and a modelling/mutagenesis study have revealed several relevant residues for drug binding such as Asp670, and among them also Trp300 and Trp666^21,22^ that were previously identified as essential in rescue experiments of the SV2A KO^11^. Further, LEV was shown to reproduce some of the deficits in neurotransmission seen in SV2A/B DKO when incubated in wild type neurons over an extended period and after trains of neuronal stimulation^23,24^. Together, these data make it plausible that LEV interferes with the function of SV2A and that the binding of LEV could localize to a substrate binding pocket, as supported by the functionally essential residues Trp300 and Trp666 in the TM-part of SV2A. To settle questions regarding LEV’s binding site and obtain further insight into the nature of the presumed substrate binding pocket of SV2A, high-resolution structures would be required.

SV2 proteins also constitute protein receptors for botulinum neurotoxins (BoNTs)^25–27^ together with tetanus neurotoxin (TeNT), the most potent biological toxins known^28^, which makes SV2 proteins clinically relevant for other reasons than their involvement in neurotransmission and epilepsy. SV2 proteins are the receptors for the BoNT serotypes A1, A2, C, D, E and F^25,29–35^ (with varying preference depending on the animal species and SV2 isoform) and presumably TeNT^26,27,36^, although for the latter case conflicting data has been published^37,38^. Tetanus has been a major burden for the largest part of human history and remains a cause for neonatal death in less vaccinated regions of the world, yet its mechanism of entry to central neurons, i.e. typically GABA/glycinergic interneurons (Renshaw cells) in the anterior horn of the spinal cord has remained unclear. TeNT recognizes lipidic and one or more proteinaceous receptors, like other CNTs. The binding of TeNT to gangliosides, glycolipids enriched on the neuronal surface, has been well described and structurally resolved with its isolated H_C_ (the receptor binding domain)^39–41^ or the holotoxin^42^ but the search for protein receptors is also complicated due to peculiar way of TeNT to enter neurons^43^. The toxin enters first motoneurons in the periphery, to travel subsequently by retrograde trafficking in endosomal structures to the postsynaptic endings where motoneurons receive signal input from central neurons. TeNT is then released into the synaptic cleft and enters the presynaptic central neurons, but differently from the case of motoneurons, by being endocytosed into SVs. Then the toxin unloads the toxic light chain (LC) through the translocation domain (H_N_) into the cytosol of the presynaptic terminal where synaptobrevin-2 is cleaved, which blocks neurotransmission. Recently, Nidogens, extracellular matrix proteins, were reported as receptors for TeNT at neuromuscular junctions^44^, also involving the tyrosine phosphatase LAR^45^. For central neurons, Thy-1, a GPI-anchored glycoprotein enriched in lipid rafts in neurons, was proposed as a receptor for TeNT^46^ but other data supported SV2A and SV2B as TeNT receptors^36^. These latter findings have been questioned because a preincubation of stimulated neurons with TeNT-H_C_ did not compete with binding of BoNT/A-H_C_ for neuronal entry^37^. Further, no binding to SV2 proteins was detected in this study; the authors thus concluded that TeNT enters SVs through a different receptor. However, the seemingly incompatible findings of the two studies could probably be explained if TeNT was not binding to the same site on SV2 as BoNT/A, which could be probed with full-length and ideally native SV2A protein to rule out bias from aberrant glycosylation.

We aimed to advance the molecular characterization of the essential SV-resident SV2A on a structural level and thereby address the binding of the anti-epileptic drug LEV as well as contribute to our understanding of TeNT protein receptor recognition on central neurons. Here, we report cryo-EM structures of immunoaffinity purified SV2A, in complex with TeNT, unveiling an unknown binding site on SV2A and the simultaneous binding of co-purified endogenous gangliosides by the toxin. Further, we provide a structure of LEV-bound SV2A that reveals the binding of the drug into a substrate pocket in an outward-open conformation of SV2A, involving also functionally essential residues and suggesting an explanation for LEV’s selectivity for SV2A over SV2B.

## Results

### Purification of native SV2A

To obtain SV2A protein for structural analysis by cryo-EM, i.e. the investigation of CNT binding to full length protein and the localization of the racetam site, we evaluated previously generated nanobodies (Nbs) against rat SV2A, for their potential to purify SV2A from detergent extracts of brain tissue by immunoaffinity chromatography. This method would also ensure potentially relevant native glycosylation patterns on the protein for subsequent experiments, e.g. forming complexes with CNTs. First, we converted the Nbs into Pro-Macrobodies^47^ (Suppl. Fig. 1A) to serve also as larger fiducial marker for improved classification in cryo-EM processing and included a twin-StrepTagII (TSII) at the C-terminus. Purified tagged PMbs were preloaded on Streptactin resin and incubated with mouse brain detergent extracts to bind solubilized SV2 which was co-eluted with the PMbs by application of biotin (Suppl. Fig. 1D). None of the tested Nbs bound SV2B or SV2C, but only SV2A (Suppl. Fig. 1C), thus our method provides homogeneous sample suitable for structural studies. SV2A is highly conserved across vertebrate species (Suppl. Fig 1B), which enabled us to purify SV2A from larger domestic animals (sheep brains, see Methods and Suppl. Fig. 2) and to investigate relevant molecular interactions of SV2A that are valid across mammalian species including humans (Suppl. Fig 1B, Suppl. Fig. 3). We could purify three different SV2A-PMb complexes through this approach from sheep brain which eluted in size-exclusion chromatography (SEC) as monodisperse species (Suppl. Fig 1E).

### Analysis of SV2A-PMb complexes and identification of an SV2A-PMb-TeNT-H_C_ complex

For structure determination we subjected the SV2A-PMb complexes to cryo-EM analysis and obtained maps that provided information about the recognized epitopes on SV2A (Suppl. Fig. 1F). For SV2A-PMb2 the resolution was limited to ∼5.8 Å but the SV2A-PMb1 complex was resolved to a global resolution of 4.1 Å. PMb1 and PMb2 occupied the C-terminal end of the luminal domain 4 (LD4), the site that is also bound by BoNT/A1 and BoNT/A2 (as shown for the isolated LD4 of SV2C^29,30,48,49^). For the SV2A-PMb5 complex we could initially not obtain clear 2D-classes, and could thus not localize the epitope for this Nb. We suspected that PMb5 binds to a flexible, potentially cytosolic epitope and that the LD4 is by itself a flexible element. Since PMb1 and PMb2 occupied the described sites for BoNT/A we considered the SV2A-PMb5 complex as good starting material to investigate complex formation with receptor binding domains (the non-toxic C-terminal domains of the toxin’s heavy chain, H_C_) of BoNT/A1 and TeNT. With a freely accessible LD4 in a full-length SV2A protein we would provide a sample that could be bound also in unforeseen configurations since we hypothesized that earlier conflicting data^36,37^ could be reconciled because of non-overlapping binding sites on SV2A for either BoNT/A or TeNT. When the SV2A-PMb5 complex was incubated with the H_C_ of BoNT/A1 or TeNT (TeNT-H_C_) we observed co-elution of the SV2A-PMb5 complex with TeNT-H_C_ in SEC, but not of BoNT/A1-H_C_ (Suppl. Fig 4A, B). The tripartite complex containing TeNT-H_C_ was subjected to cryo-EM and we obtained a high-resolution map for TeNT-H_C_-bound to the SV2A-PMb5 complex (Fig. 1A, Suppl. Figs. 5 and 6A). The structure of this tripartite complex could be solved to a global resolution of 3.25Å (∼3Å in the TM part) and served as the basis for our following analyses. The TeNT-H_C_ binding apparently stabilized movements in the LD4 substantially which allowed us to obtain a high-quality map. Further processing of data from the dipartite SV2A-PMb5 complex revealed that PMb5 becomes discernible also in the absence of TeNT-H_C_ albeit at low resolution and requiring long collection times, explaining why it initially escaped our analysis (Suppl. Fig. 1F). The map of the tripartite complex resolves PMb5 and TeNT-H_C_ unambiguously and shows that PMb5 binds on the C-terminal end of the LD4, similar to PMb1, whereas TeNT-H_C_ was clearly resolved on the N-terminal end (Figs. 1A, B, 3D). The occupation of the C-terminal site on the LD4 by PMb5 readily explains why we could not form a complex with the BoNT/A1-H_C_.

**Fig. 1:**
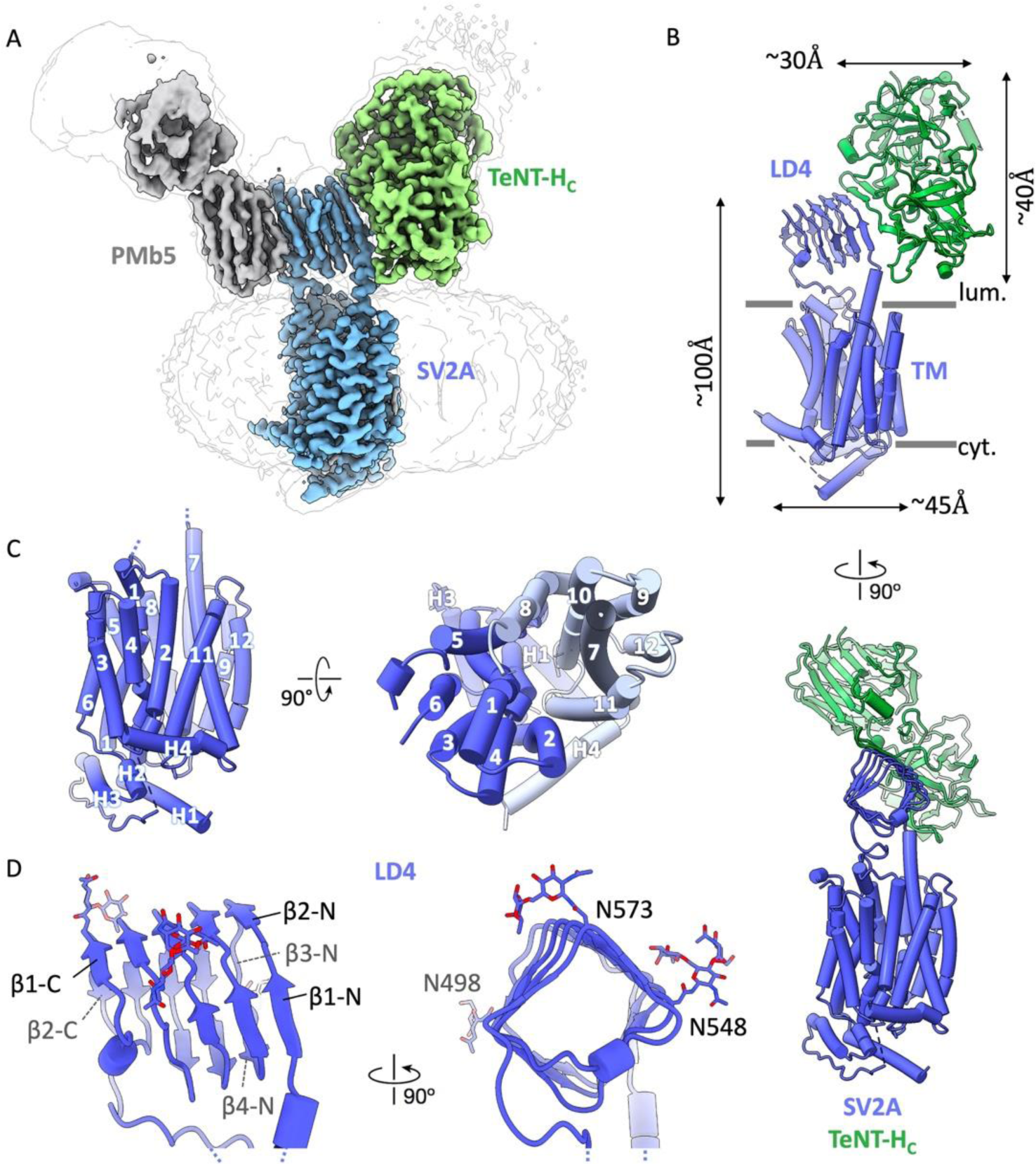
The cryo-EM structure of native sheep SV2A in a ternary complex with PMb5 and TeNT-H_C_. (**A**) Cryo-EM map of SV2A (blue) at a resolution of 3.25Å with bound PMb5 (grey) and TeNT-H_C_ (green). The micelle is outlined in light grey. The map was contoured at 0.12σ in ChimeraX (**B**) Model with dimensions of the SV2A-TeNT-H_C_ complex viewed from the sides. (**C**) Transmembrane part of SV2A with numbered helices from the side and a top view from luminal (N- and C-halves colored in blue and grey, respectively) (**D**) The SV2A-LD4 with relevant β-strands labelled and showing the N-glycans. Viewed from the side and along its longitudinal axis (from C-terminal to N-terminal).

### The structure of SV2A in the SV2A-PMb5-TeNT-H_C_ complex

The N-terminal cytosolic 137 amino acids (AA) of SV2A, a region to which Syt binds, were not resolved and may thus be natively unstructured, like the predicted fold. The rest of the protein, except for some flexible loops (AA 323-330, 401-420), could be unambiguously assigned to the map to build a model (Fig. 1A, B). SV2A features an architecture reminiscent of many MFS transporters with twelve TM helices of which the N-terminal 6 TM helices are pseudo-symmetrically related to the C-terminal half (Fig. 1C, Suppl. Fig. 7A). The SV2A-PMb5-TeNT-H_C_ complex in our sample represents likely an outward open state characterized by a cavity between the N-terminal and C-terminal halves that opens to the luminal side and is roofed by the LD4 (Fig. 2A). The cavity is lined by transmembrane (TM) helices 1,2,4 and 5 on the N-terminal half and 7,8,10 and 11 on the C-terminal half and extends to the functionally relevant residues Trp300 (TM helix 5) and Trp666 (TM helix 10)^11^ which are also involved in LEV binding^20–22^. The electrostatic potential of the cavity is negative as analysed with APBS (Fig. 2B). Whether this negative potential serves an interaction with positively charged solutes or ions, e.g. protons, requires further investigations. On the cytosolic face, SV2A features a characteristic helix (H1) before the TM part that is extended in parallel to the membrane surface and in close apposition to the lipids of the membrane. Three additional helices after TM helix 6 in a loop between the N- and C-terminal halves are arranged like a triangle with H4 positioned nearly 90° relative to H1 (Fig. 1B, C).

**Fig. 2:**
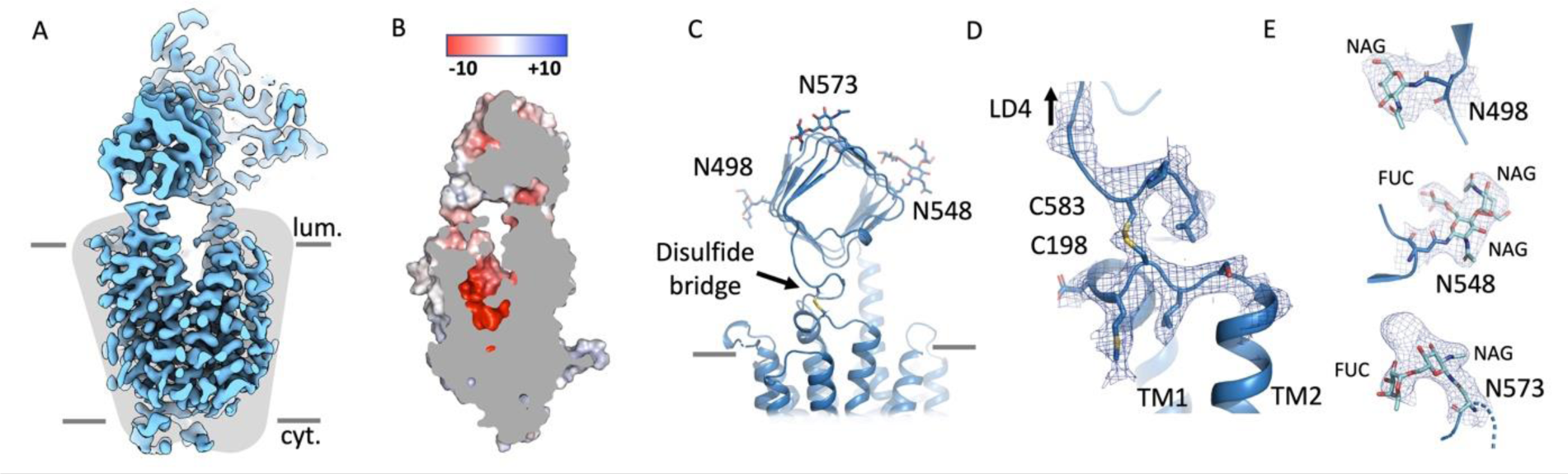
Structural features of SV2A. (**A**) Cryo-EM map of SV2A showing the outward-open conformation with the two halves (N- and C-) indicated as grey shadows. (**B**) Surface electrostatics as calculated with APBS showing an overall negative charge (red) in the transmembrane cavity between the N- and C-halves of the TM part. (**C**) The luminal SV2A-LD4 with the disulfide bridge between Cys198 (loop between TM helices 1 and 2) and Cys583 (preceding TM helix 8) and the resolved N-glycans as viewed from its C-terminal end. (**D**) Density map and model of the luminal disulfide bridge. (**E**) Density map and models of the protein-proximal ends of the N-glycans at the three N-linked glycosylation sites of SV2A - Asn498, Asn548 and Asn 573. One to two NAG units and α-1-6-glycosidic-bonded fucose are modelled. In (D) and (E) the filtered map is shown at a contour level of 9 and 8σ (Pymol), respectively.

The LD4 is a unique domain that is present in all three isoforms of SV2, but not in the closely related SV resident SVOP^50,51^. The LD4 forms a quadrilateral β-helix (Fig. 1D, Suppl. Fig. 7A) as seen in bacterial penta-peptide repeat proteins^52,53^ but in no other transporter in humans. Its structure was previously solved as isolated domain (from SV2C or as chimeric SV2C/A version^29,30,35^). The LD4 is connected to the TM part through the N-terminal helix 7 and on the C-terminal end by TM helix 8. An additional anchorage point is formed through a disulfide bridge between Cys198 after TM helix 1 and Cys583 at the C-terminal end of LD4, preceding TM helix 8 (Fig. 2C, D). Despite the multiple connections to the transmembrane part, we conclude that the LD4 is relatively flexible from the maps with different PMbs. The function of this unusual domain remains unclear, but a regulatory function could be possible as the LD4 is connected directly to the cavity-lining TM helices 7 and 8 and additionally to TM helices 1 and 2 through the disulfide bridge. The domain could thus influence four TM helices in their movements and positions and thereby impact a transport function. In the map we noted a density that reaches from the N-terminal end into the hydrophobic interior of the LD4 and that likely represents a DDM molecule (Suppl. Fig. 7B). Due to the hydrophobic interior (Phe-repeats) (Suppl. Fig. 7C) hydrophobic substances would be prone to bind there, but whether the LD4 would serve the perception of such molecules is currently difficult to state.

We noticed that SV2A is not residing in the center of the micelle (Suppl. Fig. 7D). This feature is obvious from our maps already at lower resolution and irrespective of the PMb bound or whether the TeNT-H_C_ is present. We could not find any additional density originating from a transmembrane binding partner in the extended detergent disc, but a loose association with SV2A might cause this. A mass spectrometric analysis of gel bands of copurified proteins (Suppl. Fig. 7E) did not suggest an established transmembrane binding partner, also not described interactors such as Syts were identified. It is known though that the interaction of Syt is stronger with SV2B^54^. SV2A has been mentioned as highly glycosylated protein (hence its name), although numerous other transporters in SVs may be similarly glycosylated (e.g. VGLUT2 also has three Asn-linked glycosylation sites). For SV2 proteins the glycosylation was suggested to be involved in transmitter storage^18^ and required for the binding of CNTs, as seen in glycosylated human SV2C-LD4 in complex with BoNT/A1^30^. Our preparation is natively glycosylated and allowed the identification of the protein-proximal part of three N-glycans on the LD4. An N-Acetylglucosamine (NAG) group is attached to each of the three Asn (N498, N548 and N573). A second NAG is attached to the first NAG (through β-1,4 glycosidic bonds). They are apparently fucosylated through α-1,6-glycosidic bonds to the first NAG unit at the Asn (Figs. 1D, and 2C, E) which has also been reported recently in a mass-spectrometric analysis of SV2A, except for the N-glycosylation site at Asn548 where no glycosylation was detected in the corresponding peptide^55^. We do not see indications for very large structures like keratan-sulfates, that have been proposed for SV2 proteins^2,18^, such as aberrant migration in SDS-PAGE. However, from our structural information, keratan sulfate-I could not be excluded as the protein-proximal part of these structures has an identical composition to common N-glycans. A recent mass-spectrometric analysis of SV2A refuted however the presence of keratan sulfates on SV2A^55^.

### TeNT-H_C_ binds SV2A on the N-terminal end of the LD4 by parallel β-strand augmentation

Unexpectedly, TeNT-H_C_ is bound to the N-terminal end of the LD4 domain of SV2A through an extended interface of the β-hairpin of the TeNT-H_C_ with the terminal β-strand of the LD4 in parallel arrangement by β-sheet augmentation (Fig. 3A, B). Although reminiscent of the binding mode of BoNTs A1 and A2 as seen with the isolated SV2C-LD4, TeNT-H_C_ binds to an open-ended β-strand of the LD4 (β2-N) in opposite direction and not by anti-parallel β-sheet formation seen for BoNT/A1 and A2 (Fig. 3D). While it appears in principle plausible for CNTs to bind also on the N-terminal end of the LD4, there has been no case described yet. The TeNT-H_C_ forms four hydrogen-bonds between the β-hairpin through the main-chain amines and carbonyl oxygens to extend a parallel β-sheet. Here, Ser1156_TeNT_, Tyr1157_TeNT_, Thr1158_TeNT_ and Gly1160_TeNT_ interact with the β2-N strand of LD4 (Arg491_SV2A_ and Glu493_SV2A_) (Fig. 3B)). Additional side-chain interactions are strengthening the interaction further (Glu493_SV2A_ with Tyr1157_TeNT_ and Asn1159_TeNT_, His494_SV2A_ with Asn1159_TeNT_ as well as the carbonyl oxygen of Gly489_SV2A_ and Arg1168_TeNT_ (Fig. 3B)). In addition, there is a hydrogen bond between the carbonyl oxygen of Ala482_SV2A_ in the LD4-preceding TM-helix 7 with Asn1280_TeNT_ outside of the LD4. We see no interactions with the three N-glycans on the LD4 domain that would contribute to the binding of TeNT-H_C_ to SV2A. This is in line with previous mutagenesis studies^36^ and in contrast to the glycan-H_C_ interactions described for BoNT/A1^30^.

**Fig. 3:**
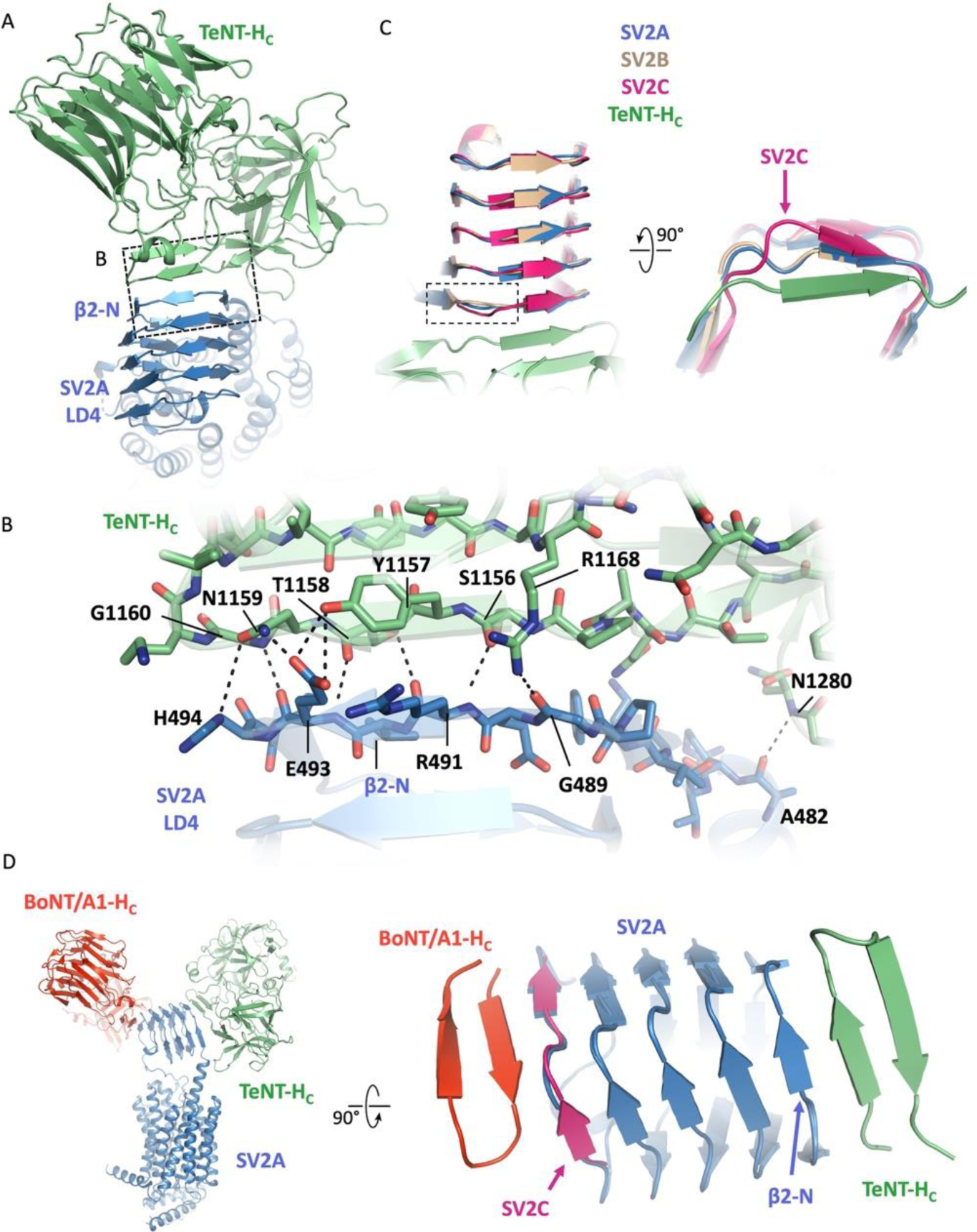
TeNT-H_C_ interactions with SV2A. (**A**) Top view from luminal on the interface of TeNT-H_C_ (green) and the SV2A LD4 (blue), showing the parallel beta-strand orientation of the TeNT-H_C_ β-hairpin bound to β2-N of the LD4. (**B**) Enlarged view of boxed area in (A) depicting residues of β2-N of SV2A-LD4 (blue) and the β-hairpin of TeNT-H_C_ (green) that contribute to the binding and the formation of a parallel β-sheet. Main chain- and side chain-interactions shown with black dashed lines. (**C**) AF2-models of the LD4 of human SV2C (pink), human SV2B (wheat) aligned with the sheep SV2A cryo-EM structure (SV2A in blue, TeNT-H_C_ in green) seen from top and turned by 90° viewed onto the N-terminal side and including the first strand of the TeNT-H_C_ β-strand. The SV2C β2-N strand shows a deviating trajectory (indicated with a pink arrow). (**D**) BoNT/A1-H_C_ (orange) in complex with SV2C-LD4 (4JRA) was aligned through the LD4 domain to the SV2A-TeNT-H_C_ complex in blue and green. The SV2C LD4 is omitted for clarity. View from top (right panel) shows the antiparallel orientation of the BoNT/A1-H_C_ β-hairpin at the C-terminus of the LD4 and the parallel β-sheet formation between the TeNT-H_C_ β-hairpin and the SV2A LD4. The last strand of the SV2C LD4 is shown in pink.

It is worth to note that the isolated recombinantly expressed LD4 of SV2C^30,48,49^ and an SV2A/C chimera^35^ are not forming the same quadrilateral arrangement of the β-strands at the N-terminus of the LD4 (β1-N is not formed, β2-N incomplete) as seen in the full-length SV2A protein (Suppl. Fig. 4C). Consequently, revealing the binding of a CNT (i.e. TeNT) to the N-terminal end of an isolated LD4 would likely not succeed; it is apparently only possible in the full-length context due to the difficulty to obtain isolated LD4 β-helix domains that contain all windings. TeNT was reported to bind SV2A and B but not SV2C^36^. Albeit no experimental high-resolution structures of SV2B and C are available, the Alphafold-2 (AF2) models of these isoforms show that the β2-N strand of the SV2C-LD4 differs significantly from SV2A and SV2B (Fig. 3C) which provides a possible explanation for the lack of SV2C interaction of TeNT and further supports that the structure of our complex represents a natural TeNT-H_C_-SV2A interface. To address that the observed binding of TeNT-H_C_ to SV2A is not a result of PMb5 binding, we subjected also an SV2A-PMb1-TeNT-H_C_ complex to cryo-EM and could obtain a map that revealed the same binding site of TeNT-H_C_ on the N-terminus of LD4 (Suppl. Fig. 4D). Supplementary Figure 4E shows an overlay of the TeNT-holoenzyme^42^ with our SV2A-TeNT-H_C_ structure that illustrates the relative positions of the translocation domain (H_N_) and the catalytic light chain (LC) towards SV2A and the membrane surface. Notably, a BoNT/A holoenzyme could also bind to a TeNT-occupied SV2A-LD4, at least in this static model.

### Co-purified complex gangliosides are bound by TeNT-H_C_ in the complex with SV2A

TeNT binds to gangliosides as nearly all CNTs and it was reported and structurally demonstrated that this occurs through two independent binding sites, the R-site (Arg1226_TeNT_) and the W-site (Trp1289_TeNT_) on the TeNT-H_C_ ^39–42^, but whether these two sites are occupied by complex gangliosides at the same time is less clear^26^. We recognized two clear densities that extend from the detergent micelle into the R- and W-sites (Fig. 4A) and could, after a focused refinement, fit the carbohydrate moieties of two gangliosides with a good level of confidence into the densities. The best fitting ganglioside for the density at the R-site is GD2 and for the W-site it is GD1a (Fig. 4B-D). GD3, GT3, GT2 and GD2 gangliosides were reported to bind into the R-site respectively, hence the GD2 ganglioside identified in our map is in line with previous results^39^. For the W-site, GM1a, GT1b and GD1a were identified as binders^40,42^, also in good agreement with our data. Notably, we have not added any gangliosides to the purified SV2A-PMb5-TeNT-H_C_ complex, the lipids are thus endogenous and stem from the sheep brain membranes. The structure supports the concept that TeNT-H_C_ has two functional binding sites for complex gangliosides, that can be simultaneously occupied which was still an open question^26^.

**Fig. 4:**
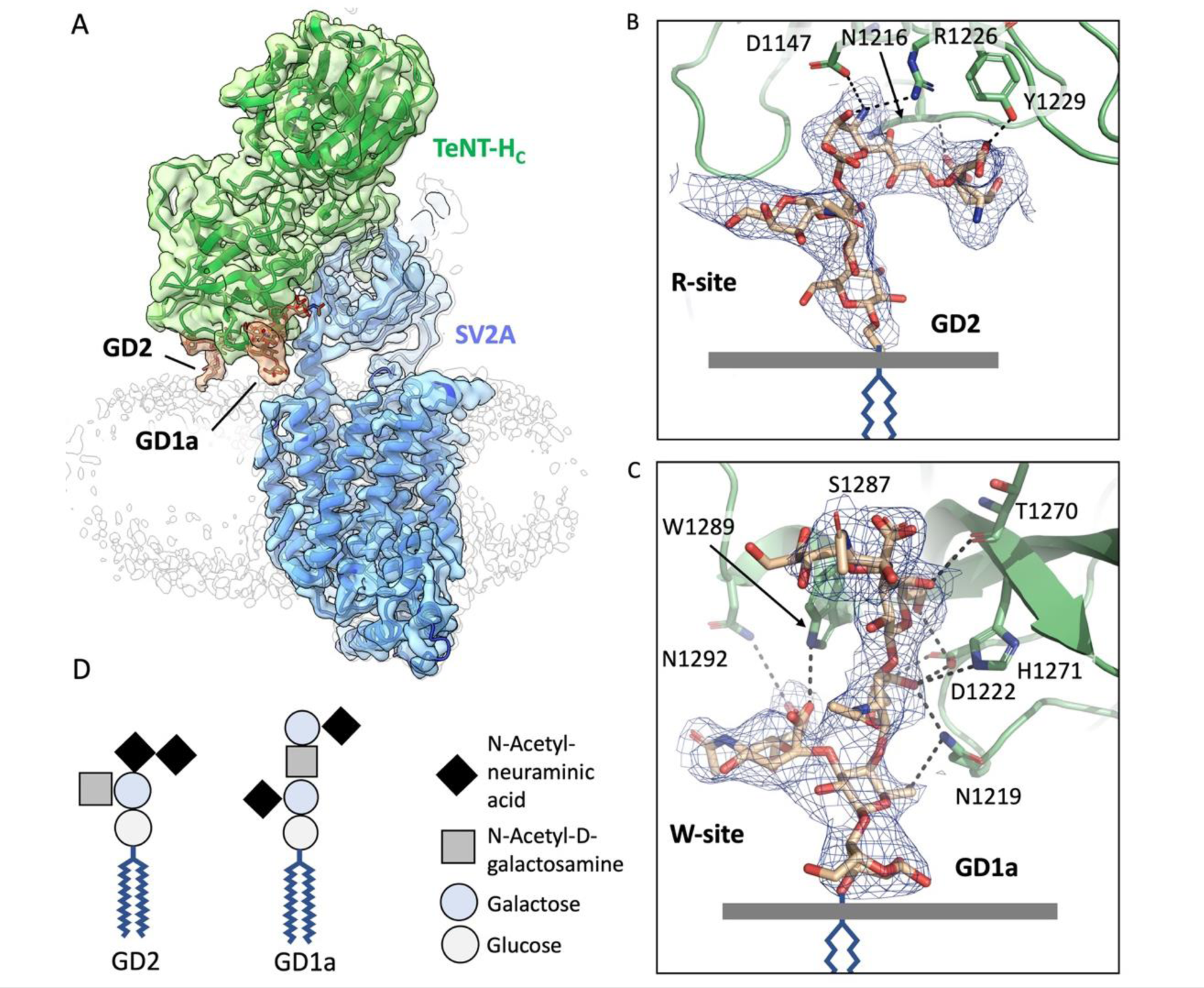
Co-purified gangliosides bound to the TeNT-H_C_ – SV2A complex. (**A**) Cryo-EM density map of TeNT-H_C_ (green) bound to SV2A (blue) and gangliosides (brown) GD2 and GD1a bound in the R- and W-sites of TeNT-H_C_ respectively. The detergent micelle is shown as outline in light grey. The map is contoured at a level of 5.4σ in ChimeraX (**B**) Enlarged view of the density in the R-site with models of GD2 ganglioside (brown) and TeNT-H_C_ (green). Residues interacting with gangliosides are shown as sticks and main interactions depicted with dashed lines. (**C**) Enlarged view of the density in the W-site with modelled GD1a ganglioside as described in (B). In (B) and (C) the density modified map is contoured at 0.8σ (**D**) Composition and assembly of GD2 and GD1a gangliosides.

### Dipartite SV2A-ganglioside receptors reveal precise angular alignment in the β-hairpin of CNTs

The H_C_-domains of the different CNTs, including TeNT, are structurally very conserved. While the largest part of the H_C_ of BoNT/A and TeNT can be almost perfectly aligned, a clear difference can be seen for the β-hairpins that bind to SV2 (Suppl. Fig. 8A). When the structures of BoNT/A1-H_C_ (or BoNT/A2-H_C_) and TeNT-H_C_ are aligned, a difference of ∼35° between the β-hairpins of a BoNT/A-H_C_ and a TeNT-H_C_ is apparent (Suppl. Fig. 8B). Importantly the β-hairpin angle is nearly independent of being bound to SV2^30,48,49,[and this study]^ or crystallized without bound SV2 moiety^34,56^, with only minor deviations, that are more pronounced for BoNT/A serotypes. The angular difference between the N-terminal β-strand of the SV2A-LD4 to which TeNT-H_C_ binds (β2-N) and the C-terminal β-strand of the LD4 to which BoNT/A1 binds (β1-C) is also 30-40° as the LD4 domain is twisted along its N- to C-terminal axis. Apparently, the difference of the β-hairpin-angle relative to the rest of the BoNT/A1-H_C_ and TeNT-H_C_ is compensating the difference in the resulting distance of the respective ganglioside binding sites to the membrane to allow for dual binding of lipids and SV2. When the SV2A-TeNT-H_C_ complex is aligned over the LD4 with a BoNT/A2-SV2C-LD4 structure, both H_C_ show almost the same distance of the ganglioside binding sites to the membrane plane (∼12Å, as measured for the W-site). Both have also a preference for the same or very similar carbohydrate moieties of gangliosides that would extend to similar degree over the membrane surface. The importance of the binding angle of an H_C_ relative to the membrane plane becomes obvious when the domains are placed on the other respective ends of the LD4. For example, if TeNT-H_C_ was bound on the C-terminal end of the LD4, the R- and W-sites would almost come to lie in the membrane (too low). In turn, if BoNT/A was bound at the N-terminal end, the ganglioside binding sites where too much elevated from the membrane surface (Suppl. Fig. 8C). Although, a certain flexibility in the β-hairpin and between the N- and C-terminal domains of the H_C_ was noticed in dependence of bound gangliosides or LD4 for BoNT/A serotypes^49,56^, the reported deviations could not account for the required shift in the angle of the H_C_-β-hairpin to bind the N- or C-terminal end of the LD4 respectively. Hence, the angle of the β-hairpin has evolved to ensure binding of the respective ends of the LD4 domain of SV2 while being bound to the carbohydrate-moiety of the gangliosides.

### Levetiracetam binds to SV2A in the transmembrane region in an outward-open conformation

Besides the binding of CNTs we aimed to obtain insights into the binding of the anti-epileptic racetams by a LEV-bound structure, as these compounds are among the most important anti-seizure medications. The binding of LEV and related racetams has been extensively studied in binding assays of SV2 proteins in membranes, both, in brain tissue and overexpressing cells. To evaluate the capacity of SV2A to bind LEV in our detergent-solubilized SV2A-PMb complexes we subjected the purified protein to drug binding using thermal shift assays with CPM dye in a real-time PCR cycler. With the SV2A-PMb5 sample we could observe a clear positive shift in the melting temperature (Tm) of SV2A by 3.7° C at a LEV concentration of 250 µM (Fig. 5A,B, Suppl. Fig. 9). A control protein, VGLUT1 bound to a PMb, did not show a shift in Tm in the presence of the LEV. Hence this complex retained its potential to bind racetams irrespective of the bound PMb5 or being detergent-solubilized. Due to the high resolution of the SV2A-PMb5-TeNT-H_C_ complex, where TeNT-H_C_ acts as an additional fiducial marker and stabilizer of motions in LD4, we continued to use this combination for further structural analysis. To localize LEV in the structure, the SV2A-PMb5-TeNT-H_C_ complex was incubated for 30 minutes on ice with 250µM LEV before plunging into liquid ethane. The resulting map at 3.53Å from these data collections revealed no obvious larger differences compared to the LEV-free structure except for a clear density that is sandwiched between Trp300 and Trp666 matching to the size and shape of LEV. A focused refinement improved the resolution from 3.53 to 3.26Å, and a subsequent density modification step in Phenix refined the map further (Fig. 5D, Suppl. Figs. 6B and 10). From the side, in a sliced view, LEV locates in a negatively charged pocket that lies in the middle of the membrane (Fig. 5C). The space in the outward open conformation from the pocket towards the luminal entry stays unperturbed and accessible for a molecule with the size of LEV. LEV itself is bound to SV2A through multiple interactions, most obvious by sandwiching (π-π interaction) between the conserved Trp-residues 300 and 666 (Fig. 5E). The oxo-group of the pyrrolidone-ring (a γ-lactam) of LEV forms a hydrogen bond with Tyr462 and the branched butyl-amide moiety of LEV is positioned between TM-helices 4,5,7 and 10. The ethyl-group of this moiety points towards TM helix 4 and interacts with Ile273 through a hydrophobic interaction. In SV2B, the Ile273 position is occupied by leucine (but it is also isoleucine in SV2C) (Suppl. Fig. 11). While LEV fits snuggly into the pocket with an isoleucine at this position, a leucine would clash with the compound. This could be one important difference to explain why the SV2B isoform is not binding LEV. The amide of the butyl-amide points towards Asp670 on TM helix 7 and Cys297 on TM helix 5. The NH-group is tightly coordinated by the sidechain carboxy-group of Asp670 (∼2 Å) and the oxygen of the butyl-amide interacts with Cys297 through a hydrogen bond (2.8 Å distance) (Fig. 5E). The corresponding residue of Cys297 is a glycine in human, mouse or sheep SV2B (Suppl. Fig. 11) and could thus not coordinate the oxygen of the amide in LEV, another difference that contributes to the lack of LEV-binding in SV2B. This circumstance speaks also for the chosen placement of the amide- and the ethyl-groups of the butyl-amide moiety of LEV. We thus consider a hydrogen-bond between the LEV-amide group and Cys297 as an essential component in the binding of LEV to SV2A besides the interactions with Asp670 and Tyr462.

**Fig. 5:**
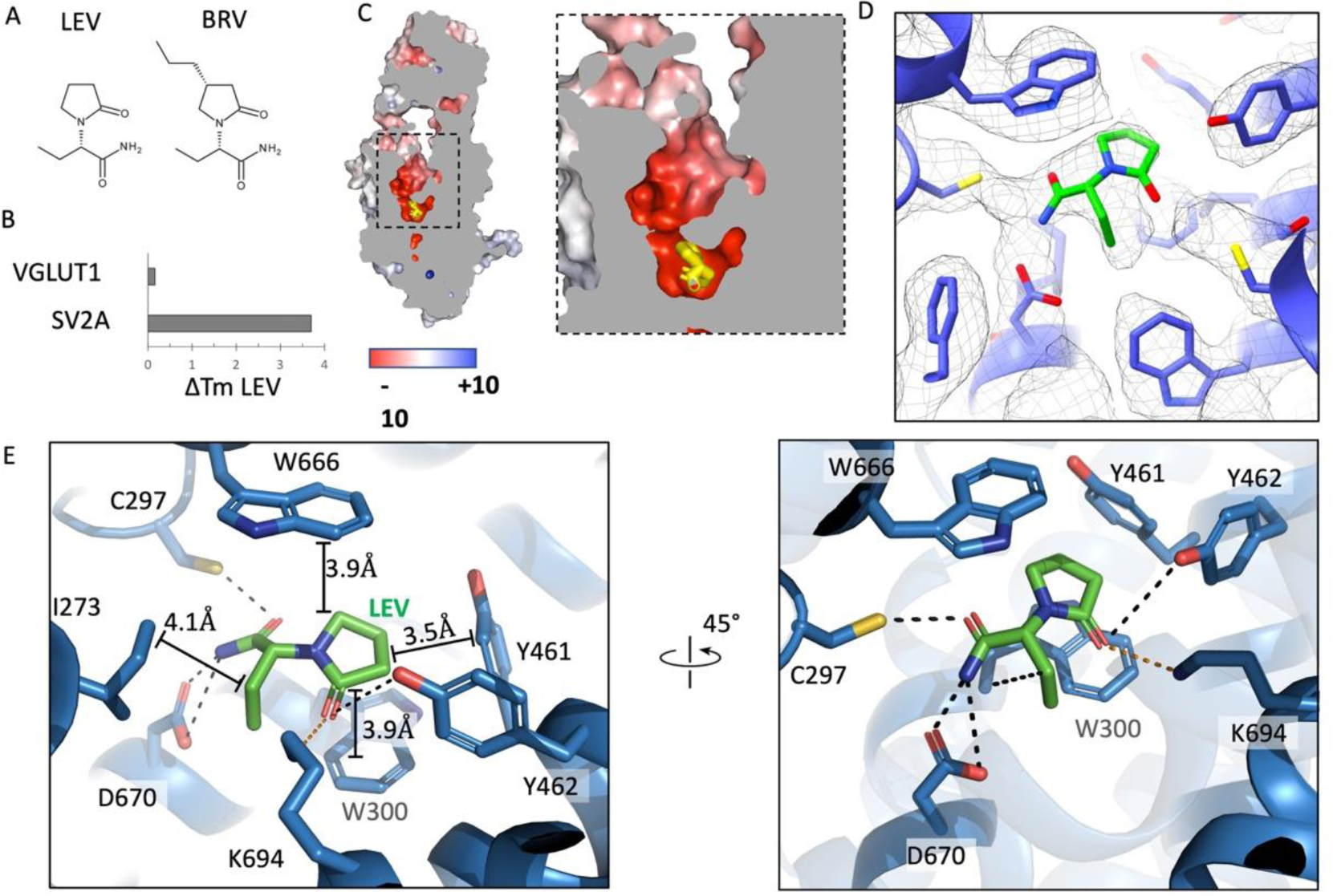
The Levetiracetam binding site in SV2A. (**A**) Chemical structure of levetiracetam (LEV) and Brivaracetam (BRV) (**B**) Elevation of the melting temperature (Tm) of SV2A in the presence of 250 µM LEV. The melting temperature of a control protein (rVGLUT1) is not affected by LEV. (**C**) Sliced side view with electrostatic surface charge and bound LEV (yellow). The boxed region is enlarged in the right panel. (**D**) Cryo-EM density map of bound LEV (green) and the coordinating residues of SV2A as sticks. The density modified map is contoured at 0.58σ in ChimeraX (**E**) Detailed view of the LEV (green) interacting residues in SV2A. Polar interactions are shown as black dashed lines. The interaction of K694 with the oxo-group of the pyrrolidone ring shown as orange dashed line due to its relatively long distance of 4.1Å. Distances of π-π interactions or other hydrophobic interactions are indicated along solid black lines. The right panel shows a rotated view from a different angle.

Brivaracetam (BRV), a more potently binding racetam differs from LEV by an additional hydrophobic element, a propyl group, attached to the pyrrolidone-ring (Fig. 5A). When assuming a similar coordination in SV2A for BRV as for LEV (an overall very similar molecule and specific for SV2A too) the propyl-group could mediate interactions within a hydrophobic patch that is built from Ile663 and Val608 and the aromatic ring of Tyr461 which explains its stronger binding. From our analysis we could not find an obvious explanation why SV2C is not binding to LEV as reported^19^ because all relevant residues for LEV binding in SV2A and SV2C are identical. It could thus be that the reason for this difference is lying outside of this binding pocket, e.g. in the path towards the binding pocket which could be obscured by bulky residues or repelling charges in SV2C or that the protein adopts a different conformation in binding assays. Finally, it could also be that the overexpression of SV2C in COS-7 cells as used for the binding assays^19^ is more problematic, in comparison to SV2A and SV2B, leading to the conclusion that SV2C is not binding LEV.

## Discussion

The SV2 family of proteins, discovered more than 30 years ago has remained enigmatic with respect to many aspects of its function. Their apparently indispensable involvement in the neurotransmission of vertebrates even exacerbates this state of knowledge. However, SV2 proteins turned out to be medically of high relevance independent of their unclear role in neurotransmission. They were unveiled as the protein receptors of many different serotypes of botulinum neurotoxins and proposed, but also questioned, to represent the receptor for tetanus neurotoxin in central (inhibitory) neurons.

Remarkably, a major drug against epilepsy – Levetiracetam - discovered in the 1970’s by UCB/Belgium turned out to bind to SV2A. Other than one might assume, the compound did not immediately uncover more of SV2’s secrets, however it would remain an important tool on the way to a deeper understanding of SV2 proteins besides its medical use.

Structural studies on CNT-SV2-interactions are founded on isolated SV2C-LD4 or SV2C-SV2A LD4 chimeras which provides us only with a limited window for probing these important interactions, and further largely on the prototypical BoNT/A. To gain more insights into the molecular details of SV2 proteins and their interactions with toxins and anti-epileptic drugs, experimental structures are needed. To provide a structural basis for the characterization of SV2A we purified the protein to homogeneity from sheep brain and subjected it to cryo-EM for the identification of the LEV binding site and further to investigate the controversial binding of TeNT to SV2A.

Our study revealed that TeNT binds to the N-terminal end of the LD4 domain in parallel orientation, a surprising finding which has not been anticipated earlier and which is unique for CNTs as for now. The structure corroborates earlier data that showed a binding of TeNT to SV2 proteins^36^ and provides strong support for their role as TeNT protein-receptors in central neurons. Further, the conflicting finding that TeNT and BoNT/A H_C_ domains do not compete for SV2 binding^37^ can be readily explained from the revealed binding site of TeNT on the opposite end of the LD4 which could apparently have both toxin H_C_ domains bound at the same time. The interface between SV2A and TeNT-H_C_, as concluded from the comparison of LD4 crystal structures and our cryo-EM structure, could only be formed with full-length SV2A due to the completeness of the N-terminal end of the LD4 β-helix. This points to potential limitations to use isolated LD4 in capturing the diversity of CNT interactions. Moreover, we could reveal the binding of the lipidic ganglioside receptors of TeNT-H_C_ while being bound to SV2A, visualizing, and reinforcing the decades old theory of dual-receptor recognition for neuronal toxin entry^57^. Whether the co-purification of endogenous gangliosides is due to interactions with SV2A cannot be answered from our data as we do not see any density proximal to SV2A TM helices that could stem from the ceramide moiety of the gangliosides. To be directly bound to SV2A, the gangliosides are also located too far away from the membrane-facing helices of SV2A. But it could be that gangliosides are enriched in the proximity of SV2A. Gangliosides have been shown to be associated with Syts^58^, hence SV2 proteins and Syts could form ganglioside rich patches that recruit CNTs with higher probability. The simultaneously bound ganglioside- and SV2A-receptors pinpoint to a scenario where the plane in which a toxin diffuses on the membrane prior to interactions with SV2 proteins, is very critical. We conclude that there is an adaption through the angle of the binding β-hairpin relative to the rest of the toxin-H_C_ to fulfil both conditions optimally, binding to gangliosides and SV2. Through this, the chance for endocytosis into SVs at extremely low concentrations of the toxin would be greatly enhanced. The structure further carefully suggests why SV2C is not bound by TeNT as the N-terminus of the SV2C-LD4 deviates substantially from SV2A and SV2B in AF2 predictions, however this requires insight into experimental structural data on SV2C for verification. Presumably, the inability to bind SV2C would ensure that TeNT is only rarely entering recycling SVs of motoneurons (mainly SV2C) but does so with great efficacy only after reaching central neurons (mainly SV2A) by retrograde transport as part of the strategy to reach the type of neurons where the toxin exerts its strongest effect.

Additionally, the newly revealed binding site for TeNT has implications for medical and biotechnological applications of CNTs. CNTs (mainly BoNT/A serotypes) are utilized in the treatment of spasms and chronic pain apart from their cosmetic uses. Immune responses can arise due to repeated application of the toxins, which could be overcome by higher potencies of the toxin. From our structure it could be concluded that BoNTs could be engineered to generate a mixed form of the toxin that targets either ends of an SV2 LD4 and chimeric versions have been generated already^59^. This should result in increased chance for entry into neurons and thus require less of the applied toxin to reduce immune responses. CNTs have also been engineered for targeted delivery of different proteins to neurons, by exchanging the light chain with a different cargo^60–62^. By using both ends of an LD4, in principle two different cargos could be delivered to neurons to name just one of the potential applications. Future engineering approaches with the H_C_ of CNTs will shed light on the possibility to employ the TeNT binding site for the abovementioned uses.

The MFS architecture of SV2 proteins has not been challenged in general, but due to the unclear function, structural insights as provided in our study certainly reinforce the concept of a membrane transporter. The pseudosymmetric 2×6 TM architecture and the luminal open cavity in an outward-open conformation resemble other MFS transporters with vesicular localization. The presumed substrate pocket, as delineated by the bound LEV molecule in the center of the membrane plane is built from highly conserved amino acids in support of their functional significance. Two tryptophan residues that sandwich also the planar LEV molecule have been shown to be essential in rescue experiments, but it remains to be seen if and which solute they would probably bind. The pocket is further featuring several polar residues in the vicinity of the central area between Trp300 and Trp666 suggesting an interaction with a substance that has at least polar elements. Recently, it was shown that SV2A would transport galactose when expressed in yeast cells. A hexose could indeed fit into the pocket and interactions with Trp-sidechains are common in carbohydrate interactions. Whether this or other sugars are transported substances, needs to be tested in SVs though. Although the function of the LD4 is currently unclear and make SV2 very unique among 12-TM MFS transporters, its connection to several TM-helices that are involved in LEV binding suggest that it is either mechanistically involved in a transport cycle or of a regulatory function e.g. through binding of yet unknown modulators. These could be either extracellular proteins, peptides or small solutes. Whether the uncapped β-strands on the LD4 serve a particular required function (i.e. the uncapped nature is part of their exact function) and are thus consequently targeted by CNTs will only become clear if we understand more about this domain.

The binding of LEV as we see it in the structure is largely in line with earlier binding studies using radiolabelled racetams^21^. These binding studies together with our structure further show that the core elements of binding in racetams are the pyrrolidone-ring and the butyl-amide group, and that the latter confers selectivity to SV2A binding. Additional groups, as for instance the propyl-group in BRV increase binding strength but would not relocate the compound to a different pocket. The resolution of our structure allowed us to identify LEV unambiguously in the pocket. However, the butyl-amide moiety branches into two groups, the amide and the ethyl-group which are of similar size. We reconstructed the most plausible rotamer by taking the chemistry and information from the SV2B isoform into account. With this in mind, the placement of the ethyl-group towards Ile273 becomes more likely than pointing to Cys297 for two reasons. First, Ile273 is a leucine in SV2B and would clash with the ethyl-group, in line with the fact that SV2B does not bind LEV. Second, if the ethyl-group was pointing towards Cys297, it would be facing a glycine in SV2B. While a glycine residue would not be very much interacting with the ethyl-group, it would also not be prohibitive and thus not explain well why SV2B does not bind LEV. In LEV, the oxygen of the amide can be positioned such that it interacts with Cys297, and this still allows for the interaction of the NH_2_-group with Asp670. In SV2B, the interaction would be limited to Asp670 through the NH_2_-group, which could weaken the overall binding. While our data can explain the lack of LEV binding in SV2B, we cannot exclude that SV2C is binding LEV, because all residues in the binding pocket are identical. This could be of importance for the use of LEV as anti-epileptic drug and potentially have an effect on a smaller subset of neurons in the CNS. Whether these effects are adverse or not is difficult to say, but the binding of LEV and derivatives such as BRV to SV2C should be re-evaluated to settle this question. The structure of SV2A bound to LEV presented herein confirms earlier binding studies^21^ and opens up possibilities for the development of even more potent molecules on a rational basis.

SV2 proteins have been suggested to contribute to resetting Ca^2+^ levels in the presynaptic terminal after action potentials through uptake into SVs. The structure reveals a net negative charge in the cavity, probably attracting cationic substances or ions, however the architecture in the pocket revealed only a single acidic residue (Asp670), not sufficient for the coordination of Ca^2+^ as seen in canonical and untypical binding sites for this ion in membrane proteins^63^. Further, recent uptake experiments with radioisotopes of Ca^2+^ into SVs including SV2A/B DKO could also exclude this idea^8^. Additional efforts will be required to understand how SV2A affects neurotransmission for which this structure will be helpful in developing experimental strategies. SV2A is expressed in inhibitory (mainly GABAergic) and excitatory (glutamatergic) neurons - how the action of LEV can be understood in these two different types of neurons remains unclear until the contribution of SV2A in the cycle of SVs is understood on a molecular level. The location of LEV in SV2A supports the idea that the drug is interfering with its function as it is binding in the core of a typical MFS-fold where substrates are bound.

During the writing of this manuscript, another research group has provided a structure of SV2A with bound LEV^64^. We note that the LEV molecule was placed differently from our study with respect to the rotamer of the butyl-amide as the authors placed the ethyl-group pointing to Cys297. Higher resolution structures will be needed alongside with mutagenesis studies to resolve the exact conformation of the butyl-amide group of LEV when bound to SV2A. We note further that the published structure differs with respect to the disulfide-bridge between Cys198 and Cys583 that we can clearly resolve in the map but that seems to be not formed in the protein overexpressed in insect cells^64^. This could have impact for the mobility of the LD4. Otherwise, the published structures of SV2A (PDB-IDs 8JLC and 8JLF) and the one of this work do not show major differences at a first glance.

In summary, our work provides a structural basis for the further characterization of SV2 proteins, based on natively purified protein, and including a conserved luminal disulfide bridge as well as the native glycosylation pattern. We could reveal the precise location of the LEV binding pocket which appears to coincide with a substrate pocket of a yet unknown substance. The dimensions and chemistry of this pocket will be aiding in limiting substrate searches and future developments for improved SV2A-targeting drugs will benefit from structure-based drug design through the presented data. Further, a structure of a ternary complex composed of SV2A, endogenous gangliosides and the H_C_ of TeNT provides compelling evidence for SV2A as the long-sought-of receptor of TeNT in central neurons. The unexpected binding mode and simultaneous binding to lipids expands our knowledge on CNT binding to its receptors, settles open controversies for TeNT and opens up new possibilities for toxin engineering to address neuronal cargo delivery and improving therapeutic applications of CNTs.

## Methods

### Expression constructs

PMbs were generated as described previously^47,65,66^ but with a C-terminal twin-strep-tag II (TSII) that was inserted into the C-terminal MBP by PCR. Briefly, Nb1-5 were PCR amplified and subcloned into the vector pBXNPHM3^65^ (Addgene #11099) and fused with a C-terminal (TSII-tagged) MBP that lacks the first six amino acids (starting at Lys7) through a di-Pro linker. The resulting expression construct contains from N- to C-terminus: a pelB leader sequence, a deca-Histidine tag, an MBP, a 3C cleavage site, a Nb, the proline-proline linker, and the truncated MBP followed by a TSII, as depicted in Suppl. Fig. 1A.

BoNT/A1-H_C_, corresponding to residues 871-1296 of the holotoxin, was synthesized (Twist biosciences) in three fragments that were fused by typeII restriction enzyme cloning with *SapI*. The expression construct was subcloned into a peT28a vector using *NcoI* (encoding the start methionin) and *NotI* (Stop codon 5’ of *NotI*). The construct encodes for an N-terminal deca-Histidine tag followed by a 3C cleavage site followed by BoNT/A1-H_C_. Between the 3C-site and BoNT/A1-HC the construct contained a *BamHI* restriction site to insert other H_C_-domains using *BamHI* and *NotI*. For the expression of TeNT-H_C_, the H_C_-domain of TeNT encoding the C-terminus of the toxin from Glu875 to Asp1315 was amplified from the plasmid pKS1 (pKS1 was a gift from Neil Fairweather (Addgene plasmid # 84079^67^ with flanking *BamHI* and *NotI* sites including a stop codon 5’ of *NotI* and exchanged for BoNT/A1-HC in the peT28a vector digested with *BamHI* and *NotI* resulting in a N-terminally deca-Histidine-tagged expression construct.

### Nanobody selection

The selection of Nbs followed published protocols^68^. Llamas were immunized with membranes overexpressing rat SV2A as antigen.

### Protein expression and purification

PMbs were expressed and purified as described earlier^47,65,66^. Briefly, MC1061 bacteria were transformed with pBXNPHM3-PMb1-5, grown at 37°C in terrific broth (TB) with 100µg/ml ampicillin until an OD_600_ of 1.2 was reached and subsequently induced with 0.024% L-arabinose. Expression was performed for 3.5 hours at 37°C. Cells were harvested by centrifugation and the resulting pellet was resuspended in 50 mM Tris-HCl pH 8, 150 mM NaCl, 0.5 mM EDTA, 10% glycerol containing protease inhibitors (Roche, cOmplete, EDTA-free Protease Inhibitor Cocktail) and frozen at -30°C for later use. BoNT/A1- and TeNT-H_C_s were expressed in the same way as described for PMbs, but instead BL21-DE3 cells were transformed with the respective peT28a expression vectors under selection of 50µg/ml kanamycin and IPTG was applied at a concentration of 0.5 mM to induce protein expression at an OD_600_ of 0.6. The cells expressing BoNT/A1-H_C_ and TeNT-H_C_ were harvested by centrifugation and the respective pellets were frozen in a buffer containing 50 mM Tris-HCl pH 8, 150 mM NaCl, 0.5 mM EDTA, 10% glycerol and protease inhibitors and stored at -30°C.

For the purification of PMbs, frozen pellets were thawed, additional protease inhibitors were added and the cells were incubated with lysozyme for 30 min at RT under stirring. All subsequent steps were performed at 4-8°C. Subsequently, DNase I at 50 µg/ml and MgSO4 (5mM final concentration) were added, and cells were lysed by sonication. Cell debris was removed by ultracentrifugation in a 45Ti rotor (Beckman) at 42k rpm for 30 min. The resulting supernatant was applied to NiNTA-resin in batch for 1.5 hours, washed with 150 mM NaCl, 40 mM imidazole pH7.6, 10% glycerol and bound proteins were eluted using 150 mM NaCl, 300 mM imidazole pH 7.6 and 10% glycerol. An over-night cleavage using HRV 3C protease was performed during dialysis against 150 mM NaCl, 10 mM Hepes-NaOH, 20 mM imidazole, pH 7.6, 10% glycerol. The purification tag (N-terminal deca-His-MBP) was removed by re-binding to NiNTA resin and the flow-through containing PMbs was concentrated and subjected to gel filtration chromatography on a Superdex 200pg 16/600 column (Cytiva) which was previously equilibrated with 150 mM NaCl, 10 mM Hepes-NaOH 7.6. Peak fractions were pooled, mixed with 10% glycerol, aliquoted, frozen in liquid nitrogen and stored at -80°C.

BoNT/A1- and TeNT-H_C_s were purified in similar manner with the exception that eluted proteins at a concentration of 11 and 17 mg/ml for BoNT/A1-H_C_ and TeNT-H_C_, respectively, were dialyzed against 200 mM NaCl, 20 mM Hepes-NaOH pH 7.6, 10% glycerol without cleaving off purification tags by 3C protease cleavage. The dialyzed proteins at 11 and 17 mg/ml for BoNT/A1-H_C_ and TeNT-H_C_ were frozen using liquid Nitrogen and stored at -80°C and tested for monodispersity by injection into an HPLC on a Superdex 200 Increase 5/150 column before mixing with SV2A.

### SVA purification from sheep brain

Sheep brains from lambs were obtained from freshly slaughtered animals at the Abattoir of Brussels (Ropsy Chaudronstraat 24, 1070 Brussels, Belgium), where lambs are slaughtered two days/week for human consumption, through an appointment with the veterinarians. The brains were immediately placed on ice and kept cool during transport back to the laboratory where they were immediately processed. Sheep brains were roughly cleaned from larger blood vessels, excess connective tissue and bone pieces, cut into ∼2 cm blocks, frozen in liquid nitrogen and stored at -80°C until use. The brains were cut such that all parts of the brain were included in a round of purification (e.g. ½ brain including cortex and cerebellum) to avoid variations across different purifications in the yield of SV2A. For a typical purification, a total of two brains (∼180 g) were homogenized with a glass-Teflon homogenizer in 50 mM Hepes pH 7.6, 250 mM NaCl, 10% glycerol and protease inhibitors (Roche, cOmplete, EDTA-free Protease Inhibitor Cocktail). Subsequently, membranes were solubilized and proteins extracted by addition of (final concentrations 2% DDM and 0.2 % cholesteryl-hemisuccinate (CHS) for two hours at 4°C under constant stirring. The crude lysate was centrifuged at 42k rpm in a 45Ti rotor (Beckman) for 45 min to remove insoluble parts. The supernatant was incubated in batch with streptactin resin (IBA, Göttingen) which was pre-loaded with ∼2mg purified TS-II tagged SV2A-specific PMbs. After an incubation time of 2 hours unbound proteins were washed away with a buffer containing 30 mM Hepes-Na pH 7.6, 150 mM NaCl, 10% glycerol, 0.03% DDM and 0.006% CHS and subsequently eluted with the same buffer containing additionally 10 mM biotin. The eluate was concentrated to 6 mg/ml and frozen in liquid nitrogen for later use in differential scanning fluorometry, mass spectrometry or cryo-EM experiments. When we used the membranes of a S1 fraction through subcellular fractionation, removing already substantial amounts of plasma, cytosol and mitochondria we had slightly lower yields and SV2A preparations of comparable purity. The purification from whole-brain homogenate was therefore faster and more efficient.

### Expression and purification of VGLUT1

Rat VGLUT1 which was used as a control for thermal unfolding experiments was expressed in tsA201 cells by transient transfection and purified as described earlier^69,70^.

### Thermal shift assay / differential scanning fluorimetry (DSF)

Protein stability was assessed by means of thermal unfolding, carried out through a temperature ramp using a real-time PCR cycler (CFX connect, Bio-Rad) following previously published protocols^71^. In this assay, initially buried cysteines become accessible through thermal unfolding and react with the CPM fluorophore (7-Diethylamino-3-(4’-Maleimidylphenyl)-4-Methylcoumarin) which creates fluorescence adducts. CPM dye at 5 mg/ml in DMSO was diluted to 0.1 mg/ml with assay buffer (10 mM Hepes-NaOH pH 7.5, 150 mM NaCl, 0.03% DDM) and incubated in the dark under mild agitation for one hour at RT.

For each thermal shift assay experiment 2-3 µg of SV2A-PMb5 or rVGLUT1 bound to a PMb were diluted in assay buffer and mixed with either LEV at 250 µM or assay buffer in a total volume of 20 µl. CPM equilibrated in assay buffer was added to a final concentration of 0.02 mg/ml and incubated for 10 min before applying the temperature ramp in the PCR cycler. After an initial incubation of 90 s at 18°C, the temperature was raised by 1°C per 12 s until a temperature of 90°C was reached. The fluorescence change was monitored using a filter for FAM fluorescence. Unfolding spectra were analyzed using the CFX Maestro software, and the melting temperature Tm was extracted from the inflection point of the unfolding curves. ΔTm was calculated as the difference of melting temperatures in the presence and absence of LEV. For each condition three replicates were performed and standard deviations are indicated for the unfolding curves. The PMbs are devoid of free cysteines and remain undetected in the assay with CPM dye.

### Cryo-EM sample preparation and data collection

The SV2A-PMb5 complex at a concentration of 6 mg/ml was mixed in a 1:2 molar ratio with TeNT-H_C_ (or BoNT/A1-H_C_). Prior to complex formation SV2A-PMb5 was separately incubated with 2.5 mM maltose to restrict movements in the MBP moiety of PMb5, and TeNT-H_C_ was mixed with 0.03 % DDM and 0.006% CHS. After an incubation time of 1 hour on ice, the mixture was subjected to size exclusion chromatography using a Superdex 200 Increase 5/150 column (Cytiva) on an HPLC that was equilibrated with a buffer containing 10 mM Hepes pH 7.5, 150 mM NaCl, 0.03% DDM and 0.006% CHS. The peak fraction was concentrated to 7 mg/ml and used immediately in cryo-plunging experiments. 3ul of the complex were subsequently applied to glow-discharged holey grids (Quantifoil R1.2/1.3 Cu 300 mesh) and plunged into liquid ethane after a blotting time of 2s at 90% humidity using a CP3 plunger (Gatan). For the LEV dataset, 30 min before cryo-plunging, the tripartite complex was incubated with LEV at a final concentration of 250 µM. SV2A in complex with the different PMbs were prepared for plunging in similar manner.

Movies were collected on a JEOL cryoARM300 microscope with 300kV accelerating voltage and equipped with a K3 direct electron detector (Gatan) and energy filter with 20eV slit. 60 frames per movie were collected in automatic manner using SerialEM 4.1.0 beta26^72^ at an exposure time of 2.796 s, total dose of about 60 e^-^/Å2 and a defocus of 0.8-2.5 µm using a 3 × 3 multi-shot pattern. Movies were acquired at a nominal magnification of 60’000 and a pixel size of 0.71Å. A total of 22295 and 17809 movies were collected for the SV2A-TeNT-H_C_-PMb5 complex and LEV-SV2A-TeNT-H_C_-PMb5 dataset, respectively.

### Processing of cryo-EM data

The movies were preprocessed in CryoSPARC Live for motion correction using Patch Motion correction and the contrast transfer function (CTF) of the motion-corrected images was estimated using CryoSPARC CTF estimation^73^. All subsequent steps were done in cryoSPARC-v.4^73^. For both datasets, initially a similar approach of processing was chosen. The particles were picked using a multi-picking approach (blob, topaz^74^, template picker) and subjected to several rounds of 2D classification. Templates were generated from a small subset from blob picking for use as input during template picking. For the tripartite complex consisting of SV2A-TeNT-H_C_-PMb5, the selected particle stack after 2D classification was cleaned from duplicates and applied to multiple rounds of heterogeneous refinement using five ab-initio maps as initial input. A Non-uniform refinement was done with a stack of particles from multiple good lines of heterogeneous refinement. The resulting particle stack was then subjected to a 3D classification into 2 classes with a focused mask on the TM part. Next, duplicates were removed and the resulting stack was further processed in a non-uniform refinement. A reference based motion correction step was conducted and the polished particles were further used in a CTF refinement. A final non-uniform refinement provided a map of 3.25Å which was subsequently sharpened using DeepEMhancer^75^.

For the LEV dataset, after several rounds of 2D classification, the particles were further cleaned by multiple rounds of heterogeneous refinements in two processing lines using four and five ab initio maps, respectively. Particles from good heterogeneous refinements were combined, followed by a removal of duplicates and a non-uniform refinement. A 3D classification into 2 classes using a focused mask on the LEV binding site was performed prior to reference based motion correction. A non-uniform refinement to optimize per group CTF parameter for anisotropic magnification yielded a map at 3.53Å. The map was additionally processed using a Local refinement with a small mask at the position of LEV which improved the resolution to 3.26 Å and was further density optimized in phenix. Throughout all processing no symmetry (C1) was applied.

### Atomic model building

DeepEMhancer and Phenix-density modified maps were used for model building and for the preparation of figures. An Alphafold2^76^ model was generated for the sequences of sheep SV2A (W5QHQ4) and Nb5. 1FV2 (H_C_ fragment of tetanus toxin) and 7OMT (crystal structure of PMb21^47^) were used as initial models for model building of TeNT-H_C_ and PMb5, respectively, where for the latter the predicted Alphafold2 model of Nb5 replaced the Nb part of 7OMT. The models were rigid body fitted to the experimental cryo-EM map of SV2A-TeNT-H_C_-PMb5. Stretches in the models that were not visible in the map were removed. The model was subjected to iterative rounds of Phenix refinement (Phenix v.1.20.1-4487^77^) and manual model building in Coot 0.8.9.2^78^. For TeNT-H_C_, all amino acids could be built except for the residues 983-987. Nb5 could be built completely, while the density for the MBP moiety was not sufficient for model building, the MBP moiety of PMb21 (7OMT) was therefore rigid body fitted into the density for illustration in Fig.1A. For SV2A all residues could be modelled except for the N-terminus (1-137), and residues 322-329 and 401-420. For the model building of the LEV-SV2A-TeNT-H_C_-PMb5 complex, the SV2A-TeNT-H_C_-PMB5 model served as the starting point since the respective maps suggested very similar conformations. The model was subjected to iterative rounds of Phenix refinement and manual model building in Coot. Pymol v.2.0.7 and v.2.5.4, ChimeraX-1.3^79^ were used for the illustration of the structures and maps.

## Author contributions

S.S. and J.D.B. conceived the study and planned experiments; S.S. expressed and purified proteins, formed the complexes, prepared samples for cryo-EM, and performed DLS experiments; T.L. and J.S. performed the nanobody discovery; J.D.B. prepared grids for cryo-EM, collected and processed cryo-EM data and built the models; S.S. and J.D.B. analyzed data and wrote the manuscript.

## Acknowledgements

We thank the nanobody discovery team at the VIB-VUB Center for Structural Biology for their work and the nanobody cDNAs. We thank Dr. Yves Nerinckx (AFSCA) and his co-workers from the Abattoir of Brussels for supply with fresh lamb brains. We are grateful to Drs. Marcus Fislage (Biogenic Electron Cryo-Microscopy (BECM), Brussels) and Dirk Reiter (VIB-VUB Center for Structural Biology, Brussels) for assistance with cryo-EM data collection and technical support for workstations and IT-related questions, respectively. We thank Prof. Damya Laoui (VIB-VUB Center for Inflammation Research, Brussels) for mouse brains. We would like to acknowledge the funding provided by Vlaams Instituut voor Biotechnologie (VIB). The authors thank the members of the Brunner lab for support.

## Supplementary Information

**Suppl. Fig. 1:**
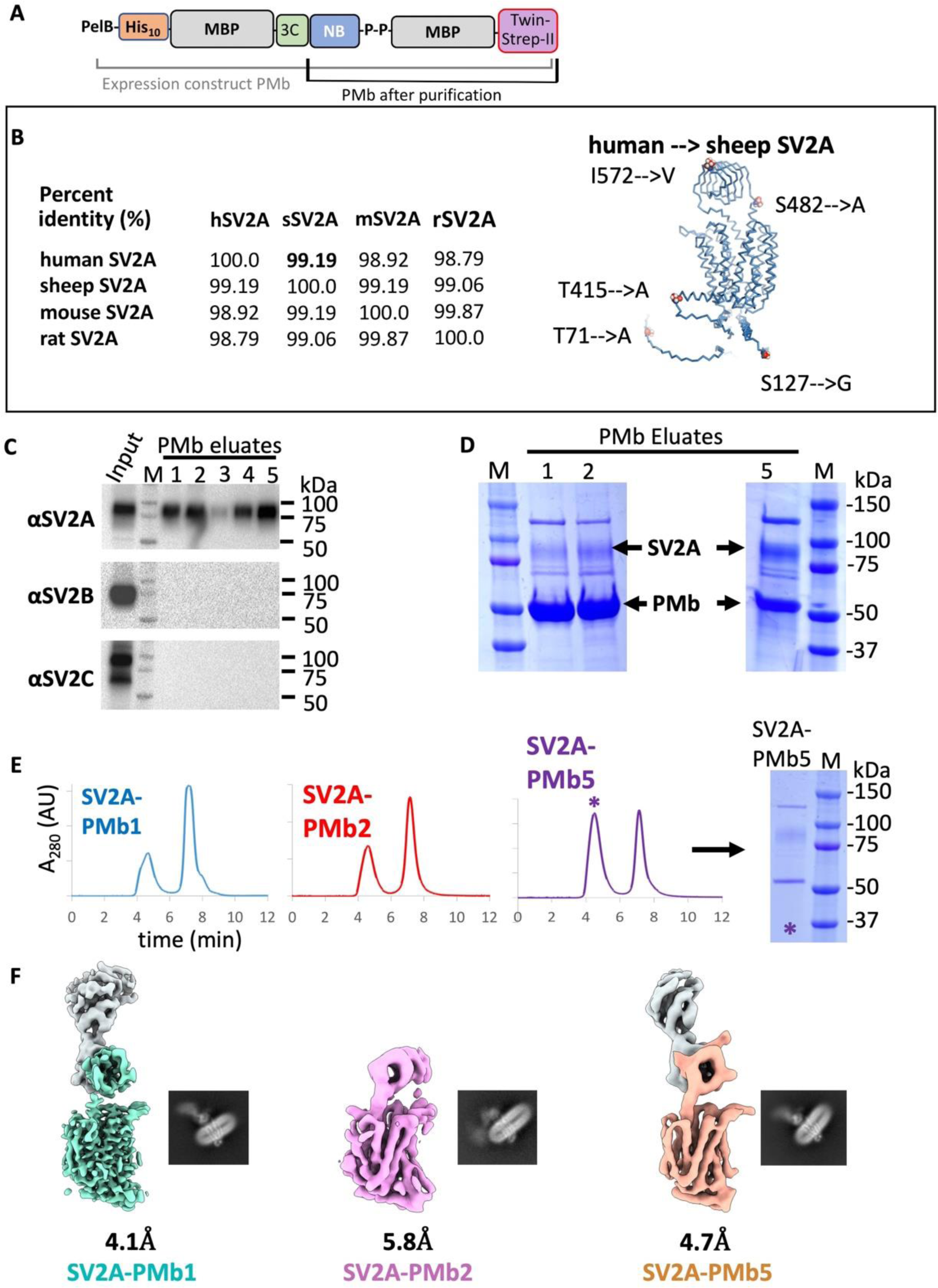
PMb constructs, PMb specificity, purification and cryo-EM maps of SV2A from brain. (**A**) Schematic representation of the PMb expression construct and final polypeptide after cleavage. pelB: pectate lyase leader sequence for periplasmic expression; 10His: deca-histidine tag; MBP: Maltose binding protein; 3C: HRV-3C-proteinase recognition and cleavage site; NB: nanobody; P-P: di-proline motif. (**B**) Identity of human, sheep, mouse and rat SV2A in percent. The amino-acid changes between human and sheep SV2A are plotted on the structure and lie all outside of the TM-region. Sheep and human SV2 have identical numbering because of identical length. (**C**) Western blot of the PMb eluates 1-5 after pull-down from mouse-brain detergent extract. The blots were probed with isoform specific antibodies. SV2B and SV2C were undetectable in the eluates. (**D**) Coomassie-stained SDS-PAGE gel of the PMb eluates with SV2A and PMbs indicated. (**E**) SEC-runs of PMb eluates 1,2 and 5 on a Superdex 200 Increase 5/150 column connected to an HPLC in 150mM NaCl, 10 mM Hepes-Na pH 7.6, 0.03% DDM, 0.006% CHS. The excess PMb elutes after ∼7.5 minutes. The SV2A-PMb complex elutes at ∼4.5 minutes. A Coomassie-stained SDS-PAGE gel with the peak fraction (marked with an asterisk) loaded is shown on the right for the SV2A-PMb5 complex with SV2A running as a smeary band at ∼80kDa and PMb5 at ∼50kDa. An additional contamination is visible at 125 kDa. (**F**) Cryo-EM density maps of sheep SV2A in complex with PMb1, PMb2 and PMb5. A representative 2D-class is shown. The maps are shown in identical orientation.

**Suppl. Fig. 2:**
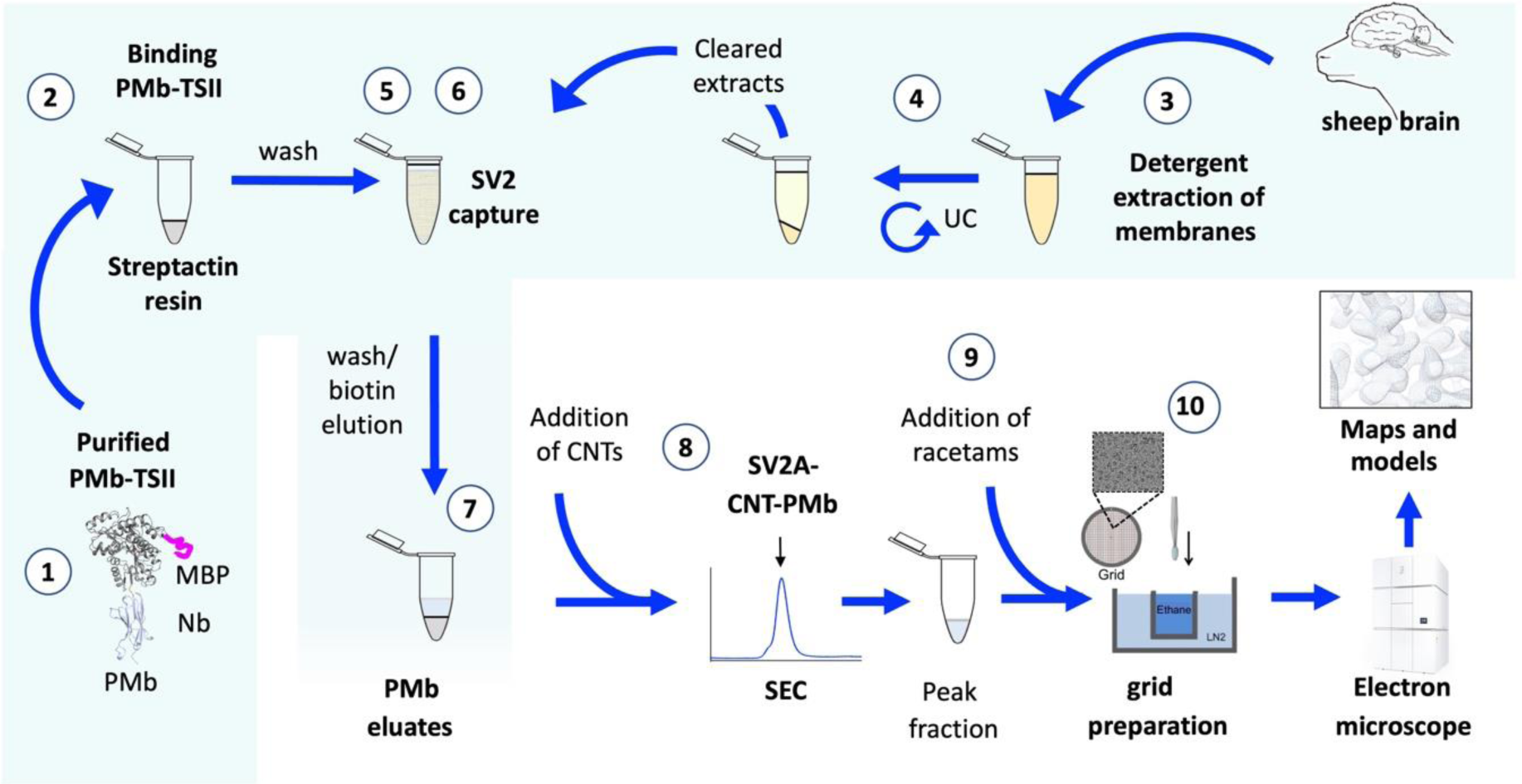
Scheme of the workflow for the purification of SV2A from sheep brain.

**Suppl. Fig. 3:**
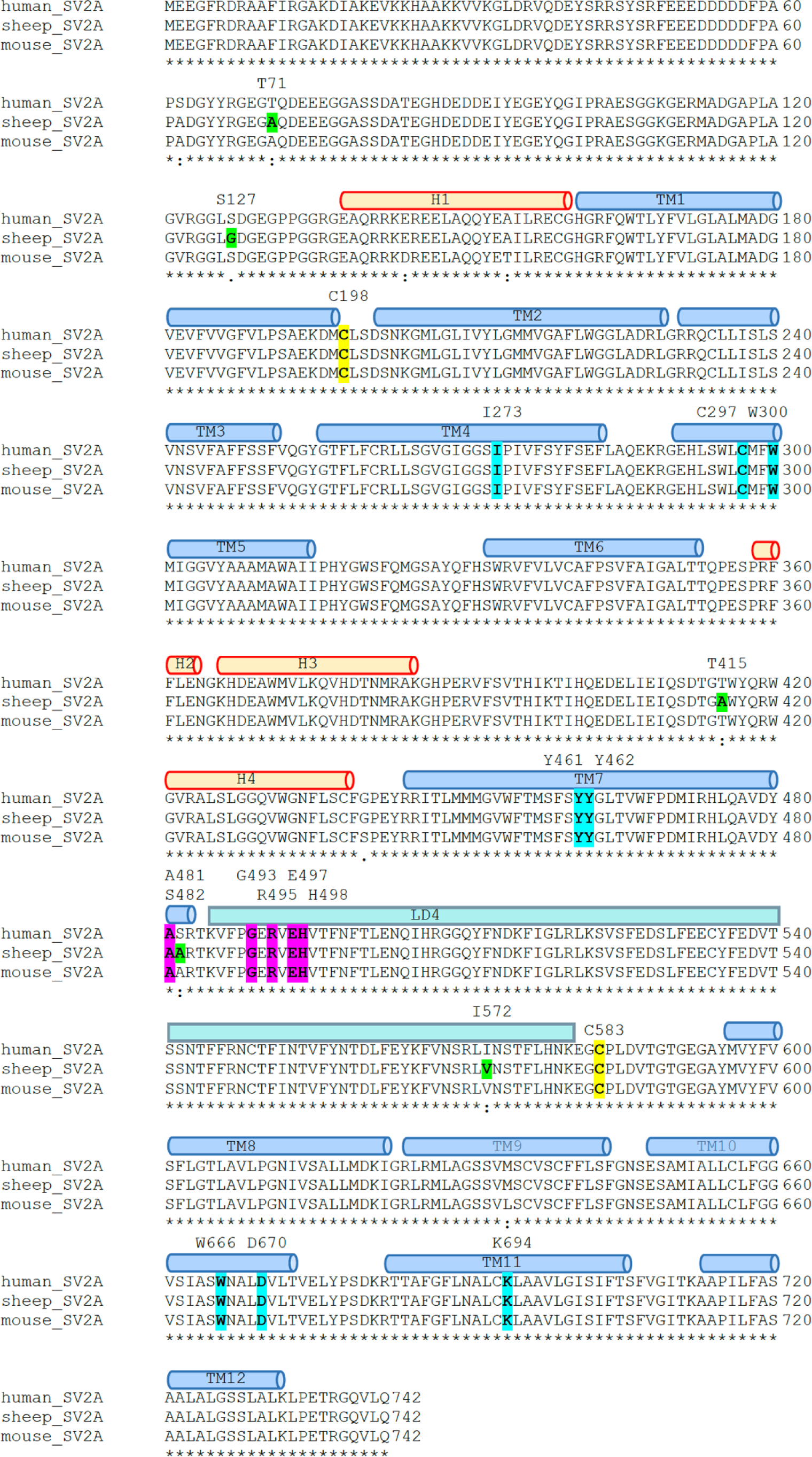
Alignment of human, sheep and mouse SV2A. Amino acid sequences of sheep, mouse and human SV2A were aligned with Clustal Omega and structural elements from the cryo-EM structure are shown as yellow cylinders (cytosolic helices), blue cylinders (transmembrane helices) and the LD4 (cyan bars). Residues that differ in sheep SV2A from the human sequence are highlighted in green. The disulfide-bonded cysteines are highlighted in yellow. Residues involved in LEV binding are shown in cyan and the TeNT binding site in pink.

**Suppl. Fig. 4:**
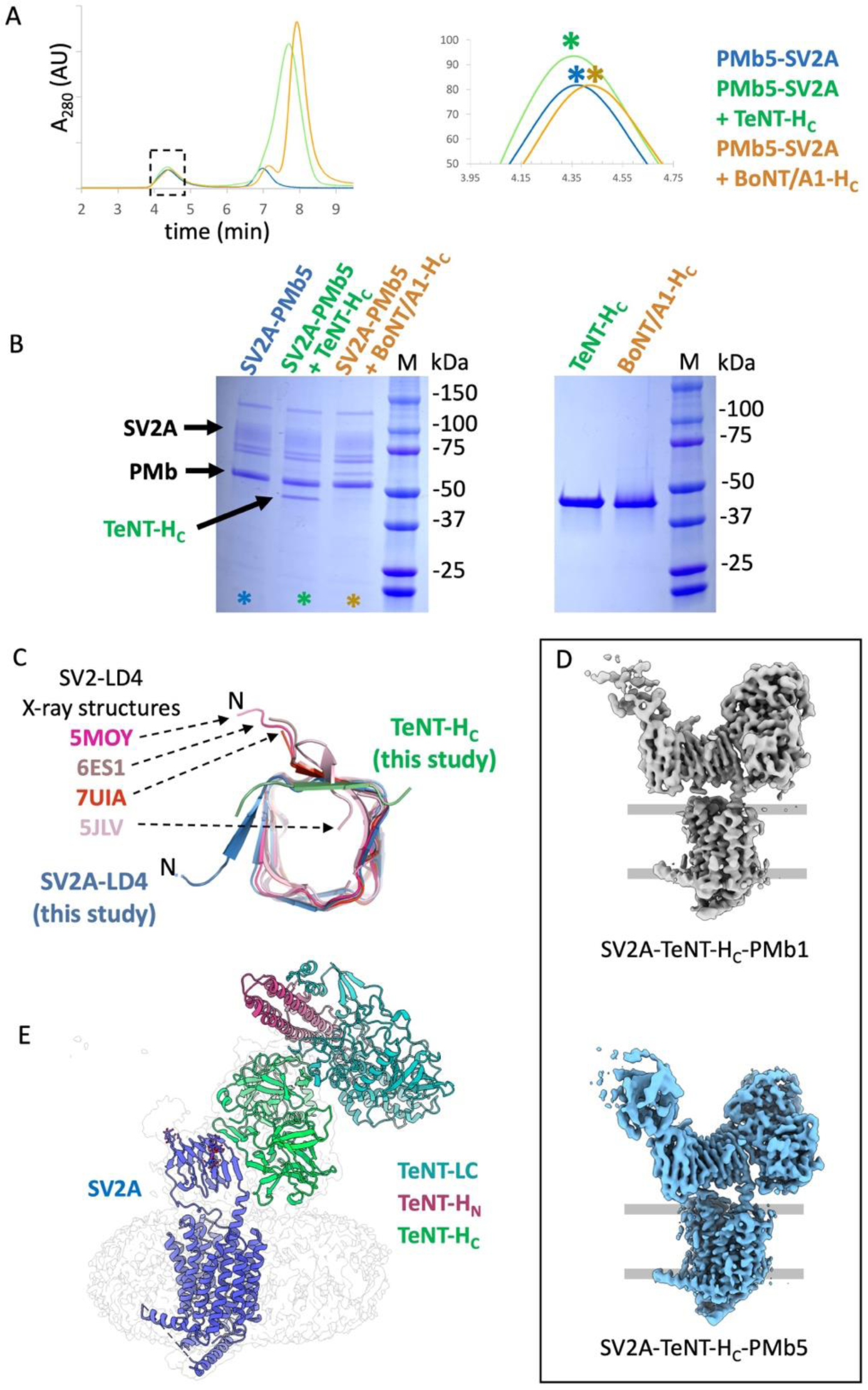
(**A**) SEC-chromatograms of SVA-PMb5, SV2A-PMb5+BoNT/A1-H_C_ and SV2A-PMb5+TeNT-H_C_ from samples injected onto a Superdex 200 Increase 150/5 column. The dashed box on the right shows the complex-peaks enlarged and marked with asterisks. (**B**) Peak fractions of (A) as marked with astrisks were loaded on an SDS-PAGE gel and stained with Coomassie blue. The TeNT-H_C_-domain co-eluted with the SV2A-PMb5 complex (TeNT-H_C_ indicated with an arrow). The gel on the right shows purified TeNT-H_C_ and BoNT/A1-H_C_ for comparison. (**C**) Crystal structures of SV2C LD4 and an SV2A/C LD4 chimera were aligned with the SV2A-TeNT-H_C_ cryo-EM structure to show the differences at the N-terminus of the isolated LD4 domains and the SV2A-LD4 domain. PDB numbers are indicated for the respective structures. (**D**) Cryo-EM density maps of SV2A-PMb1-TeNT-H_C_ and SV2A-PMb5-TeNT-H_C_. The comparison reveals the almost identical binding of TeNT-H_C_. (**E**) The SV2A-TeNT-H_C_ complex was aligned with the crystal structure of the TeNT holoenzyme (5OBN) to show the location of the translocation domain (H_N_) and the light chain (LC) and their respective position relative to the membrane plane.

**Suppl. Fig. 5:**
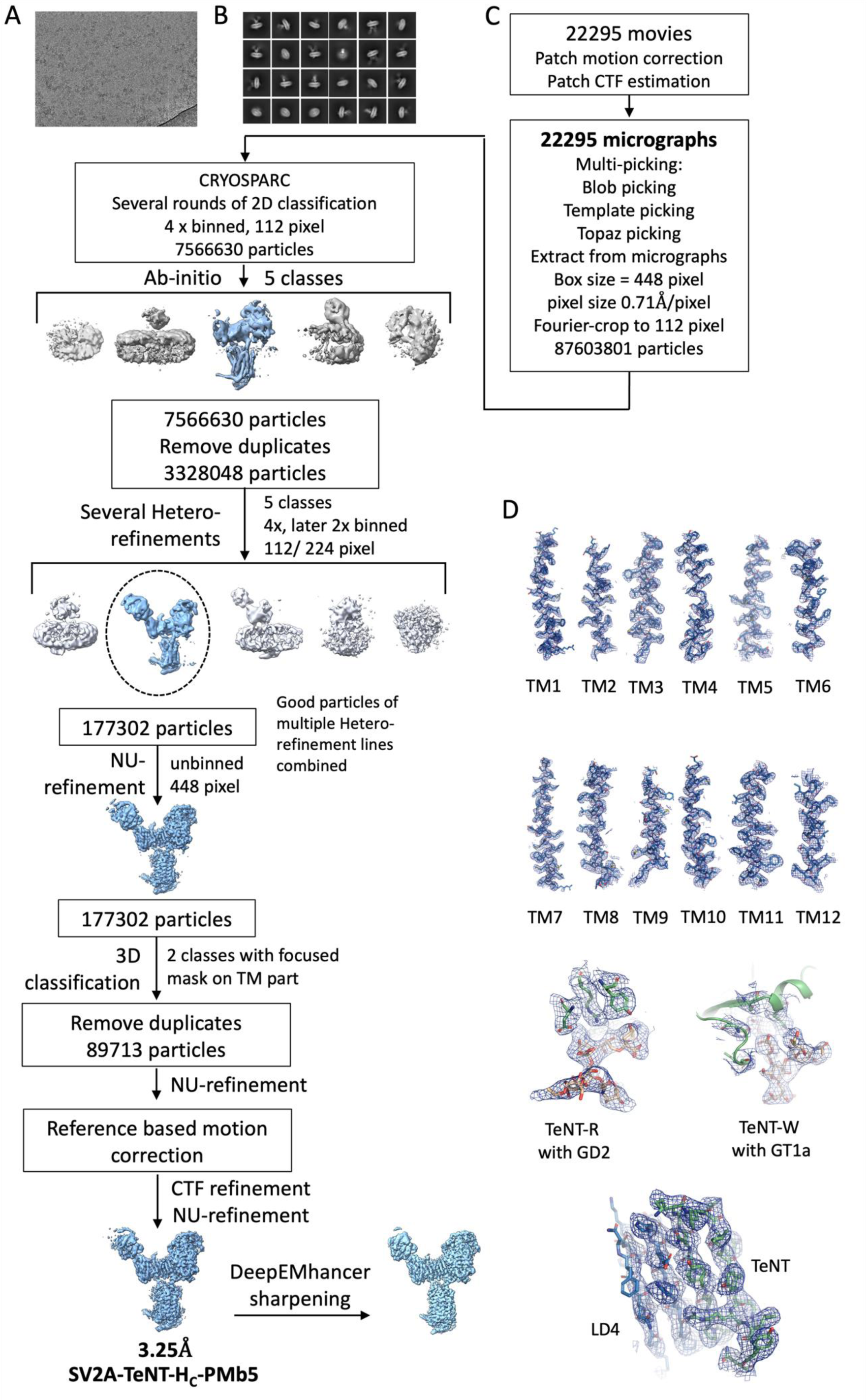
Cryo-EM workflow for the SV2A-PMb5-TeNT-H_C_ structure. (**A**) Representative micrograph from the data collection on the JEOL cryoARM300 electron microscope. (**B**) Representative 2D-class averages (**C**) Processing workflow for the cryo-EM density map. (**D**) Agreement of the cryo-EM density map with the final model shown for the TM-helices of SV2A, the densities for the gangliosides and the interface of the LD4 and TeNT-H_C_. Map contoured at 2σ in pymol.

**Suppl. Fig. 6:**
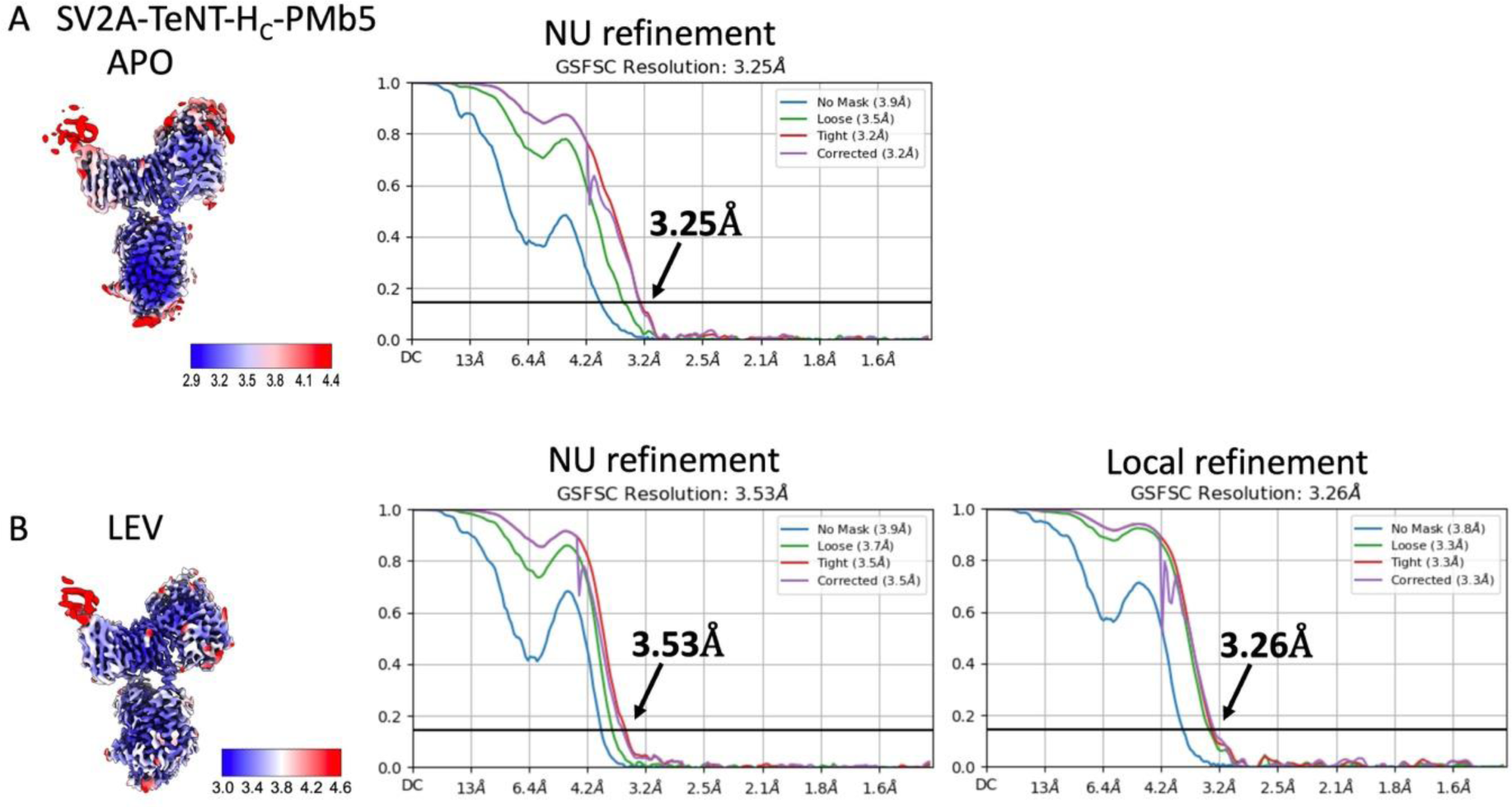
Local resolution and Fourier shell correlation plots of SV2a-PMb5-TeNT-H_C_ as Apo- or LEV-bound structures. (**A**) SV2A-PMb5-TeNT-H_C_ Apo structure local resolution and FSC plot (final NU-refinement) with the resolution marked at the 0.143 criterion. Map is contoured at 0.099 σ in ChimeraX. (**B**) SV2A-PMb5-TeNT-H_C_ LEV bound structure local resolution and FSC plot (final NU-refinement) with the resolution marked at the 0.143 criterion. The FSC plot after a final local refinement is shown on the right. Map is contoured at 0.051 σ in ChimeraX.

**Suppl. Fig. 7:**
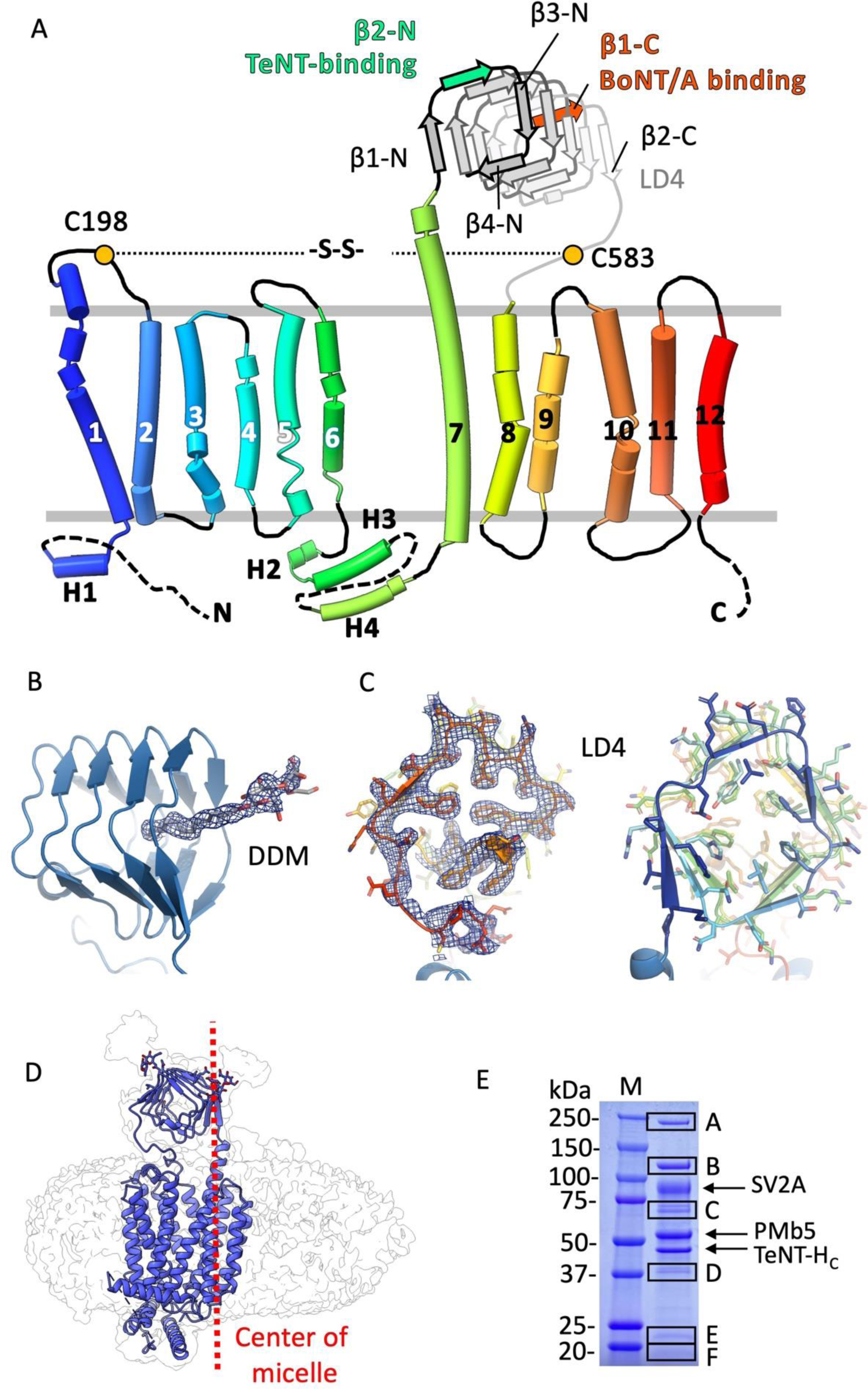
(**A**) SV2A topology coloured in rainbow from N- to C-terminus. (**B**) LD4 cartoon representation with a density located in the LD4 lumen. A DDM molecule is fitted into the density. (**C**) Cryo-EM density with cartoon model viewed from the N-terminal end of the LD4. A cartoon model of the LD4 with the same view is shown on the right to display the phenylalanine repeats pointing to the lumen. Modified density (Phenix) is contoured at 0.8σ in (B) and (C). (**D**) The position of the SV2A molecule in the DDM-CHS detergent micelle is not centered. The mass center of the micelle is indicated with a red dashed line. (**E**) Coomassie stained SDS-PAGE gel of SV2A-PMb5-TeNT-H_C_ for identification of excised bands by tryptic digests and subsequent mass spectrometric analysis. The most prominent identified proteins in the bands are: A: Acetyl-CoA-carboxylase I; B: Pyruvate-decarboxylase I; C: Methyl-crotonoyl-CoA carboxylase I; D: G-protein subunit i1; E: proteolipid protein I; F: plasmolipin.

**Suppl. Fig. 8:**
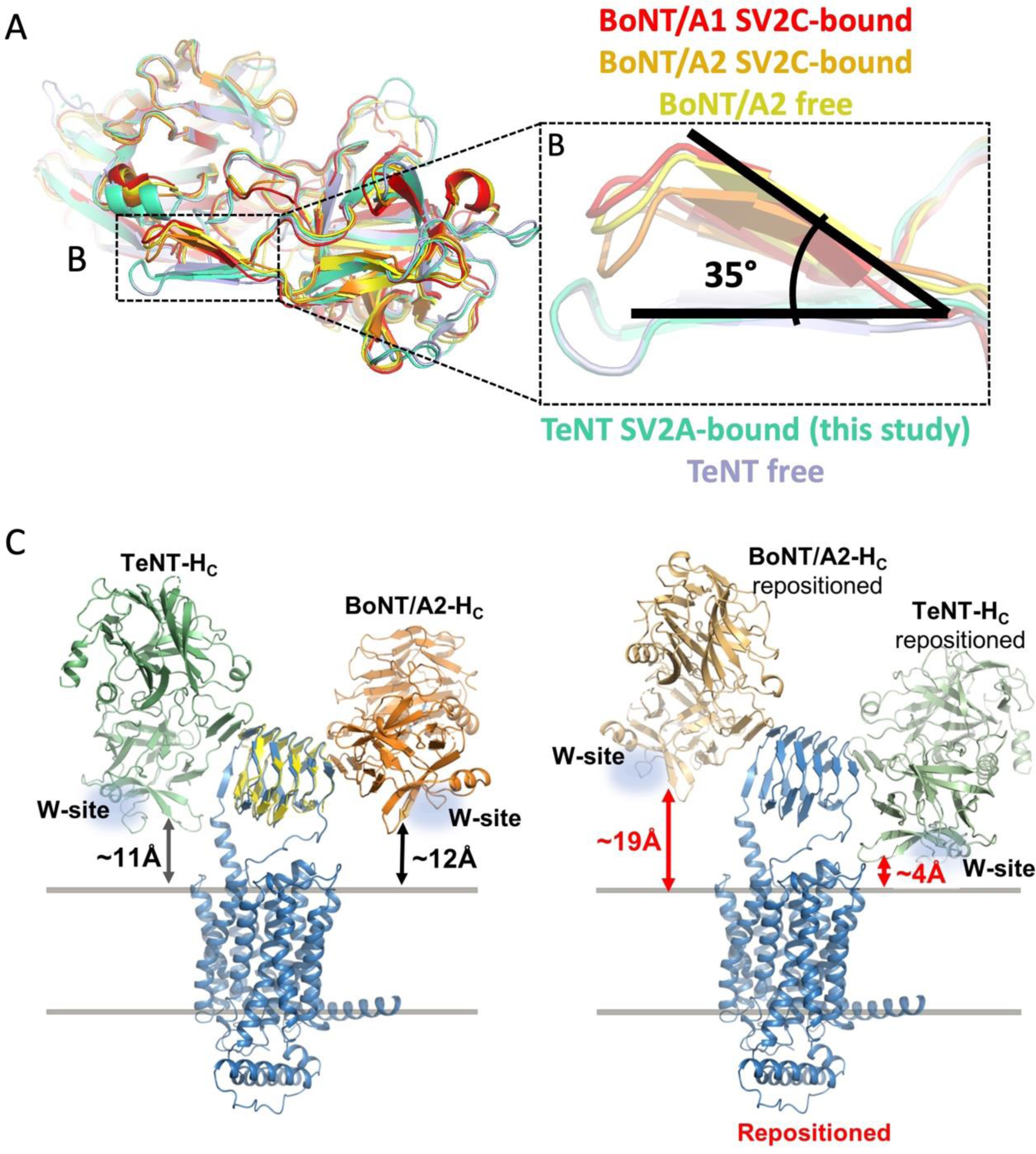
The β-hairpin angle depends on the distance of the R- and W-sites to the membrane. (**A**) Structures of LD4-bound or unbound H_C_-domains of BoNT/A1, BoNT/A2 or TeNT were superimposed. The structures align over the largest part of the polypeptide, except for the angle of the β-hairpin relative to the rest of the chain which differs by ∼35° for TeNT-H_C_ compared to BoNT/A1 or A2. The angle shows only minor variation whether an H_C_ is bound to the LD4 domains. Coordinates are from BoNT/A1-bound (5JLV), BoNT/A2-bound (6ES1), BoNT/A2-free (7Z5T), TeNT-free (1FV2), TeNT-bound (this study). (**B**) Enlarged region boxed in (A) highlighting the β-hairpin with 35° difference between the TeNT-H_C_ and the H_C_ domains of BoNT/A serotypes. (**C**) **Left panel:** The TeNT-H_C_-SV2A cryo-EM structure (SV2A blue, TeNT-H_C_ green) was superimposed with a BoNT/A2 crystal structure (orange, 5MOY) in complex with a SV2C LD4 (yellow) to align it with respect to the LD4 domain of SV2A. Distances of the last reside in the β-strand (Cα of Tyr1255 for BoNT/A2 and Gly1273 for TeNT-H_C_ within the W-site) N-terminally of the loop protruding to the membrane relative to the micelle surface where measured. Both H_C_-domains have nearly the same distance to the membrane surface plane of ∼12 Å at their native binding sites on the LD4. W-sites are indicated by a blue cloud. **Right panel:** The two H_C_ domains were exchanged and repositioned (TeNT-H_C_ to the BoNT/A2 position and vice versa). The structures were aligned on the β-hairpin of the other H_C_ domain bound to the respective LD4 terminus. When TeNT-H_C_ was positioned analogous to BoNT/A2 on the C-terminus of the LD4, the distance of the W-Site to the membrane was only 4 Å. When BoNT/A2 is positioned to the N-terminus of the LD4, the native site for TeNT-H_C_, the distance of the W-site to the membrane increases to ∼19 Å.

**Suppl. Fig. 9:**
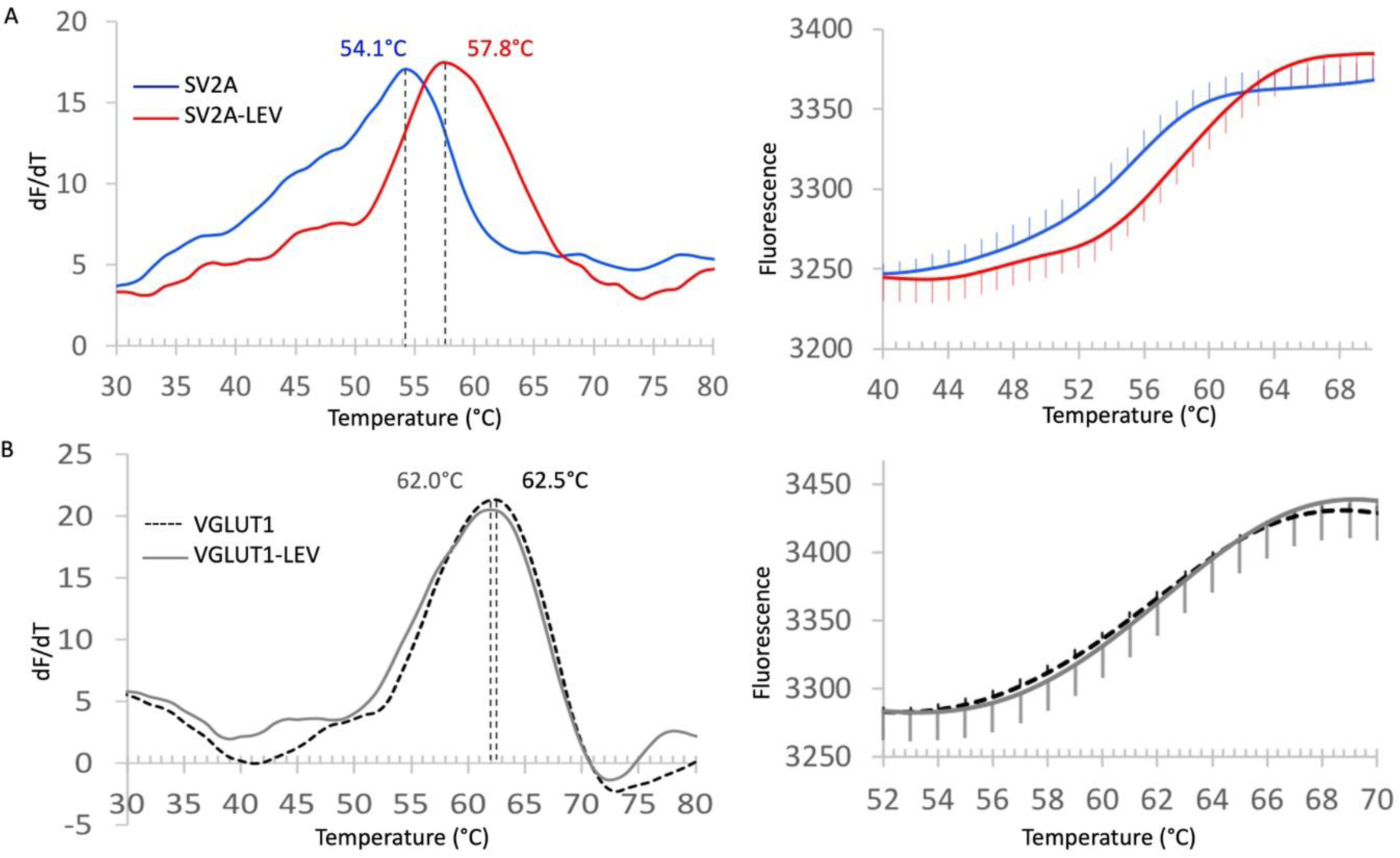
Thermal shift assay using differential scanning fluorometry to measure LEV-binding. (**A**) SV2A-PMb5 complex was heated with a ramp 1°C/12 s from 18-90° C and thermal unfolding was measured by the increase in CPM fluorescence in a real-time PCR cycler. In the presence of 250µM LEV the melting temperature (Tm) is reached at 57.8°C whereas in the absence of LEV SV2A unfolds at 54.1°C. The mean of triplicates is shown. The right panel shows the mean of the raw unfolding curve. The error bars indicate the standard deviation from triplicates. (**B**) The Tm of rVGLUT1 bound to a PMb was measured in analogous manner to SV2A in (A) in the presence or absence of 250µM LEV. The right panel shows the raw data of the unfolding curve (mean of triplicates with the error bars indicating the standard deviation).

**Suppl. Fig. 10:**
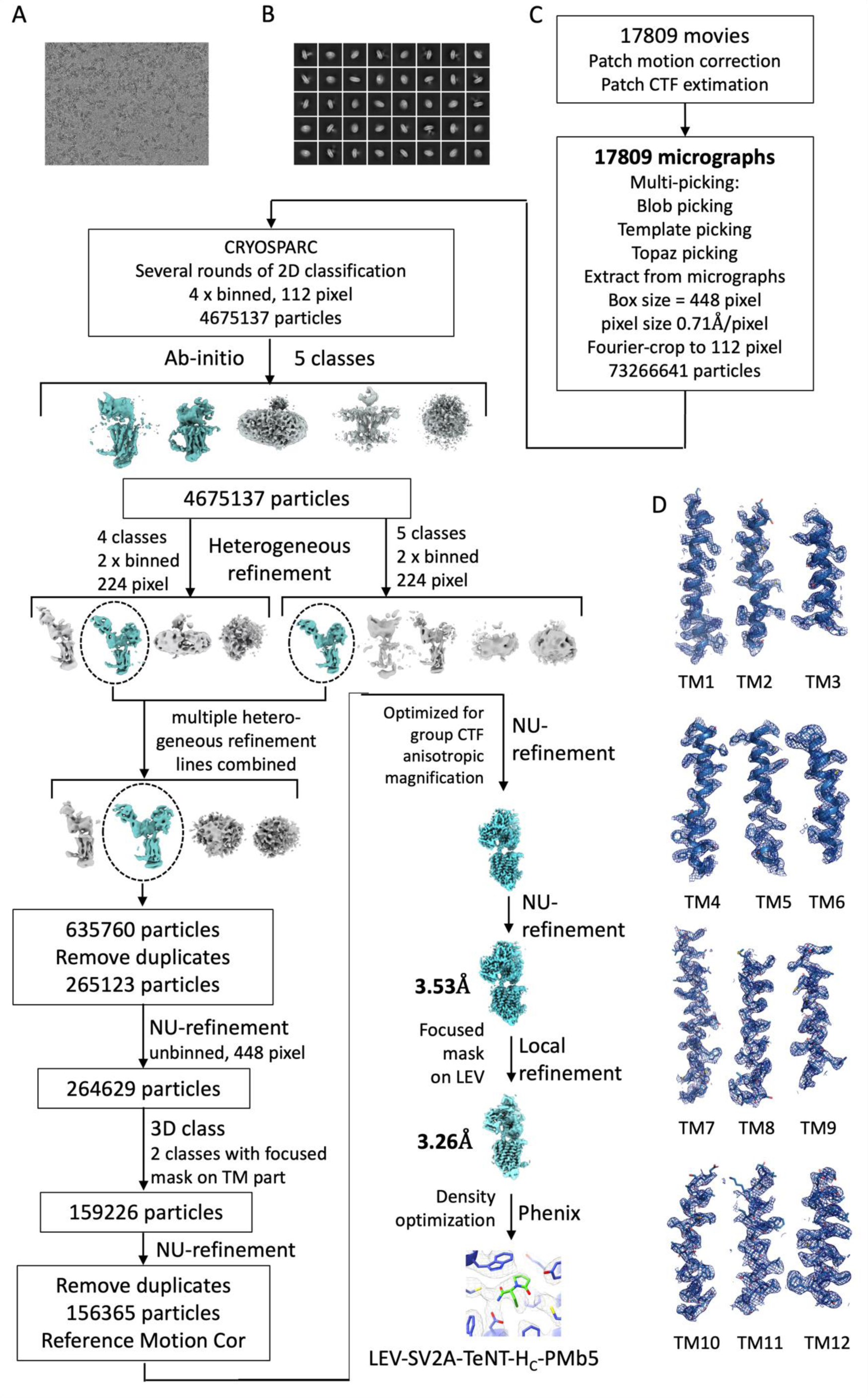
Cryo-EM workflow for the SV2A-PMb5-TeNT-H_C_ structure with LEV. (**A**) Representative micrograph from the data collection on the JEOL cryoARM-300 electron microscope. (**B**) 2D-class averages (**C**) Processing workflow for the cryo-EM density map. (**D**) Agreement of the cryo-EM density map with the final model shown for the TM-helices of SV2A. Map contoured at 2σ in pymol.

**Suppl. Fig. 11:**
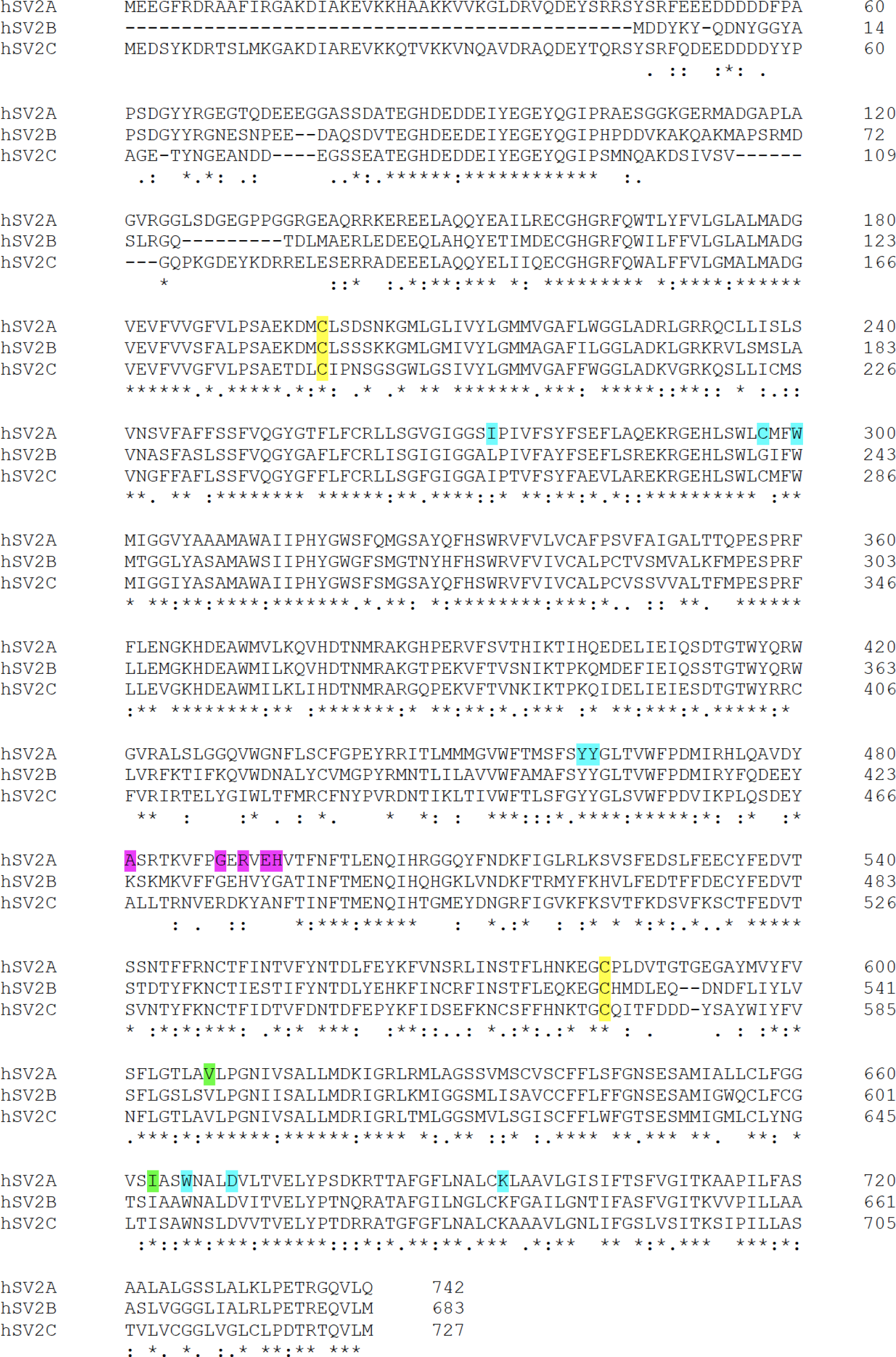
Alignment of human SV2A, SV2B and SV2C. Amino acid sequences of the three isoforms were aligned with Clustal Omega. Residues involved in LEV binding in SV2A are highlighted in cyan. Residues that are additionally important in BRV binding are highlighted in green. Residues involved in TeNT-H_C_ binding are highlighted in magenta. The disulfide-bonded cysteines are highlighted in yellow. Ile273 and Cys297 are not conserved in SV2B.

